# Hydrogel chemistry modulates mesenchymal stromal cell outgrowth and cytokine secretion profile

**DOI:** 10.64898/2026.11.26.769272

**Authors:** Wen Chen, Hao Wu, Zhengdong Gao, Hongjuan Weng, Behrad Shaghaghi, Katrien V. Bernaerts, Paul H. J. Kouwer, Paul Wieringa, Lorenzo Moroni

## Abstract

Cells are surrounded by fibrous networks that play an important role in regulating cell behavior during cell-matrix interactions. However, strategies to replicate their microenvironments with fibrous architectures are typically challenging to manipulate or even control. Here, we develop three-dimensional hybrid *in vitro* models by combining fibrous hydrogels with tunable mechanical dynamics and fibrous electrospun scaffolds. Fully synthetic polyisocyanide (PIC) hydrogels are employed to introduce tunable mechanical dynamics by simply varying the crosslinking approaches. Cells exhibited an elongated morphology in hydrogels with increasing mechanical dynamics. Moreover, a cup-shaped hollow electrospun scaffold is fabricated as a secondary fibrous component, followed by injection of hydrogels encapsulated with human mesenchymal stromal cells (hMSCs). We demonstrate that the cytokine secretion of hMSCs can be tuned by simply altering the hydrogel type or mechanical dynamics, and its functional effects highly dominate the extent of tube formation of endothelial cells cultured with different hMSC-conditioned media. These findings underscore the significance of hydrogel properties in the regulation of cell outgrowth and cytokine secretion profile, which may offer a promising approach for manipulating cell behavior via cell–matrix interactions and advancing regenerative strategies.

## 1. Introduction

Mesenchymal stromal cells (MSCs) have been demonstrated as potent mediators of tissue repair by producing and secreting a large array of bioactive factors, such as exosomes and cytokines, which can act as key regulators of tissue regeneration in regenerative medicine.[1] In addition to promote tissue repair, MSCs show immunomodulatory properties, including pro-inflammatory and anti-inflammatory abilities, which subsequently promote angiogenesis.[2, 3] Despite their outstanding therapeutic ability compared with traditional medicines, as with any cell transplantation, local cellular delivery and retention remains highly challenging. To overcome these challenges, tissue-engineered scaffolds with similar dimensional structures to those of extracellular matrix (ECM) fibres with diameters ranging from nanoscale to microscale[4] may ideally be used as building blocks to encapsulate cells. Many studies have reported the fabrication of nano- and microfibres using various techniques, such as electrospinning,[5, 6] self-assembly,[7] centrifugal spinning,[8] phase separation,[9] and drying in confined conditions[10]. Among these approaches, electrospinning has gained increasing interest due to its simplicity, versatility, and capability of producing fibrous structures that mimic the ECM, which has been demonstrated as a great potential scaffolding technique for tissue engineering applications.[5, 6] However, electrospun scaffolds often lack the three-dimensional (3D) cell infiltration, tunable mechanical properties, and bioactive environment.

Hydrogels are widely employed in regenerative medicine because they provide a highly adaptable platform for modulating cellular responses both *in vitro* and *in vivo*.[11] Their high water content, structural similarity to native ECM, and chemical flexibility allow hydrogels to create a permissive and physiologically relevant environment for cell survival, proliferation, differentiation, and functional integration.[11, 12] Importantly, biophysical cues embedded within hydrogels, such as matrix stiffness, viscoelasticity, and topography, play a crucial role in recapitulating the native cellular microenvironment.[13] These physical properties not only dictate the mechanical support provided to resident cells but also profoundly influence a broad range of cellular behaviors, including adhesion, migration, lineage commitment, and the regulation of the cell secretome.[13–15] For instance, Liu et al. demonstrated that 3D synthetic hydrogel matrices could tune stem cell secretome by varying matrix stiffness, resulting in distinct rates of wound closure in functional wound healing assays.[15] Nevertheless, how dynamic changes in matrix mechanics regulate the secretome remain unknown.

In this work, we develop a 3D *in vitro* hybrid model by combining hydrogels with tunable mechanical dynamics and fibrous electrospun scaffolds to regulate cell behavior. A hydrogel library was developed, including fully synthetic biomimetic polyisocyanide (PIC) hydrogels with similar mechanical stiffness but with different matrix dynamics (Figure 1A). These dynamics were achieved by employing physically and covalently crosslinking approaches, respectively. Natural biomaterials, including alginate and collagen were investigated as reference hydrogels. We demonstrated a direct correlation between cell morphology and the mechanical dynamics of the hydrogel matrix. Specifically, we observed that cells exhibited increased spreading in matrices with higher mechanical dynamics (Figure 1B). Moreover, a cup-shaped hollow electrospun scaffold was fabricated (Figure 1C), followed by injection of hydrogels encapsulated with human MSCs (hMSCs), resulting in the formation of a favorable 3D cellular microenvironment. Additionally, the secreted cytokines by hMSCs could be tunable through simply changing the hydrogel types and/or mechanical dynamics. The cytokine functions were investigated on endothelial cells cultured with the conditioned media from hMSCs (Figure 1D).

**Figure 1.**
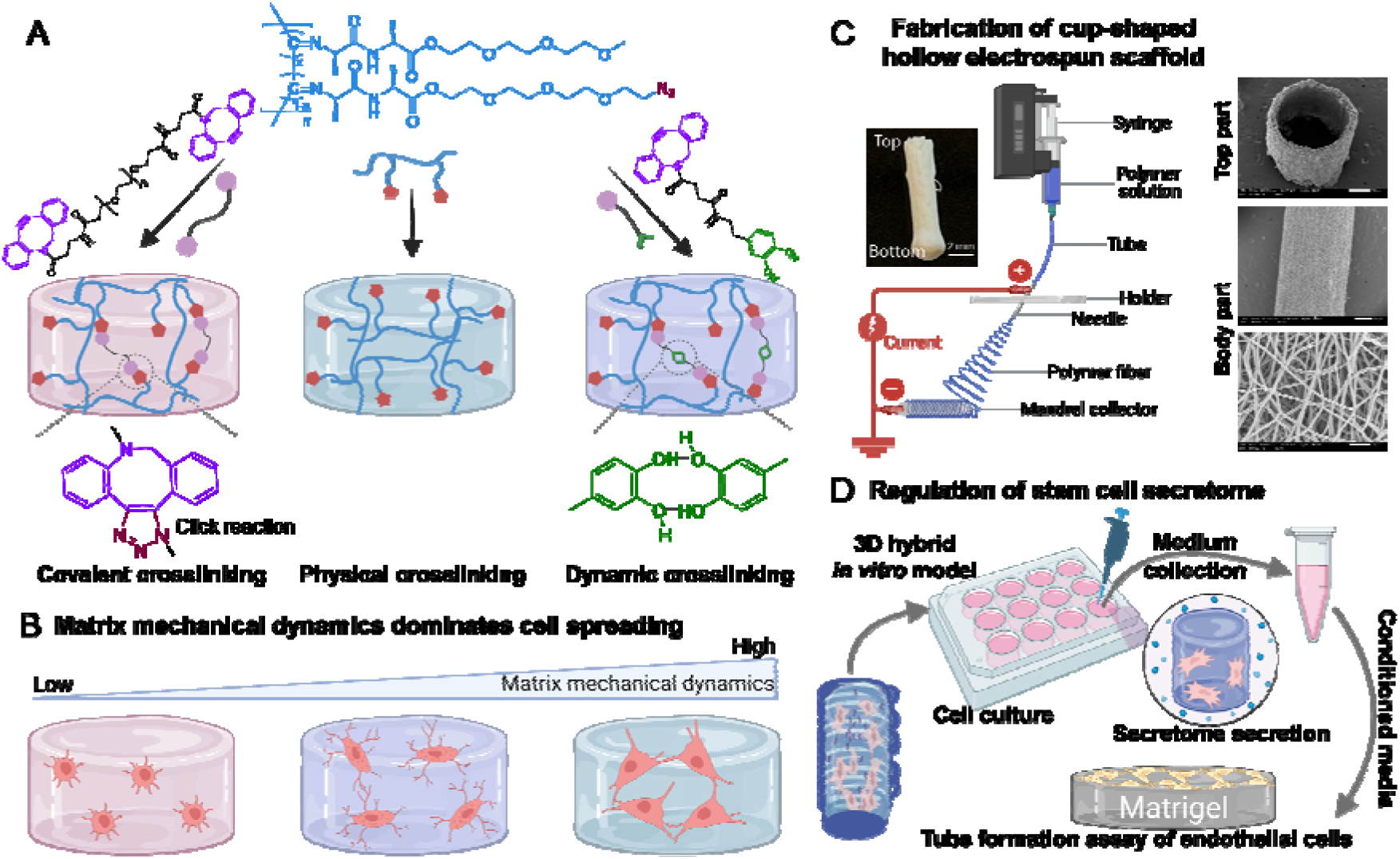
Experimental design. (A) Schematic overview of PIC hydrogels with different crosslinking strategies, including physical, dynamic and covalent crosslinking, resulting in the formation of hydrogels with tunable mechanical dynamics. (B) Cell spreading depends on the matrix mechanical dynamics. With increasing crosslinking density, cells spread less. (C) Schematic overview of fabrication of cup-shaped hollow electrospun scaffolds. SEM images show the electrospun scaffold has a fibrous structure. (D) Modulation of hMSC secretome. A 3D hybrid model is prepared by injecting cell-laden hydrogel precursor into the electrospun scaffold, followed by gelation. Stromal cells secrete cytokines during cell culture. The collected medium is used to treat endothelial cells to investigate the secretome functions by a tube formation assay. Biorender was used to draw the schemes.

## 2. Results and discussion

### 2.1 Fabrication of electrospun scaffolds

Cup-shaped electrospun scaffolds with one closed end and one open end were developed to enhance cell retention and allow cell-ladened hydrogel injection, respectively. The electrospun scaffolds were fabricated on a rounded open-end rotating mandrel with 1.8 mm of diameter, where the electrospinning needle at 30° angle was positioned towards the open-end mandrel (Figure 1C). After cutting from the mandrel, hollow scaffolds were obtained (Figure 1C, white scaffold shown in the insert image). When integrated with hydrogels via injection from the open end, we named the resulting scaffolds as hydrocups.[16] The wall thickness of the scaffolds was variable by tuning electrospinning time. However, with increasing wall thickness, hydrogels may show less integration with the electrospun scaffolds, as evidenced in previous work.[16] Therefore, we chose 120 seconds as the electrospinning time for a combination of better integration and easy handling operation. Macroscopic analysis showed that the electrospun scaffolds had microscale porous structures (Figure 1C, SEM images, Figure S1), and the wall thickness of the electrospun scaffold and the diameter of the fibers were 157.5±20.43 μm and 2.72±0.46 μm, respectively (Figure S2).

### 2.2 Preparation of a hydrogel library

The interactions between cells and their direct cellular microenvironment are interactive.[17] Cells continuously sense external forces derived from the ECM. In turn, cells remodel and reorganize the ECM, thereby allowing them to manipulate the matrix dynamics. A key characteristic of such matrices is strain stiffening, a phenomenon exhibited by natural matrices like collagen.[18] As the applied stress increases, the matrices become stiffer, a process that plays an important role in tissue adaptation by mitigating damage from structural deformations.[19] To this end, we prepared a hydrogel library to closely mimic the cellular microenvironment. This library comprises biomimetic strain-stiffening PIC hydrogels.[20, 21] PIC gels are characterized by a branched fibrous (or bundled) network that forms on heating an aqueous polymer solution beyond the gelation temperature *T*_gel_ ∼ 15-20 °C.[22] The PIC architecture and (non)linear mechanical properties show high similarities to biological hydrogels like fibrin, collagen, and actin.[18, 20] The unique structural and mechanical characteristics with high tailorability[21, 23–25] and excellent biocompatibility[21] make the PIC hydrogels as an attractive platform for wide biological applications, including *in vitro* as synthetic models[26–29], as well as *in vivo* applications for wound healing[30] and dental treatment[31, 32]. PIC gels with varying mechanics through different crosslinking approaches, including single physical crosslinking (pPIC), dual physical crosslinking (dPIC), and chemical crosslinking (cPIC) were prepared (Figure 1A). Collagen and alginate hydrogels were employed as positive and negative controls, respectively, when considering cell spreading as a metric.

PIC hydrogels were equipped with azide groups (13 mol%) based on established protocols with minor modification (see details in methods). PIC hydrogels with single physical crosslinking were prepared upon heating an aqueous cold PIC polymer solution above its gelation temperature *T*_gel_, resulting in PIC chain aggregation into bundles, thus forming a thermo-responsive hydrogel network through hydrogen bonding, analogous to the structure of hydrogels formed by intermediate filaments or F-actin. Moreover, we designed a bifunctional spacer[24] with a dibenzocyclooctyne (DBCO) group at one terminus to react with azide groups in PIC backbones through the strain-promoted azide–alkyne cycloaddition (SPAAC) cross-linking reaction, and a dopamine-based motif containing catechol groups at the other to form hydrogen bonding and/or π–π stacking between catechol groups, contributing to the secondary physically crosslinking. To prepare chemically crosslinked PIC hydrogels, DBCO-PEG4-DBCO crosslinkers were employed to form SPAAC crosslinking between azide groups in PIC polymers and each head of the crosslinkers. Additionally, collagen hydrogels were prepared by neutralization and incubated at 37°C to allow gelation, while alginate hydrogels were fabricated by ionic crosslinking with calcium ions.

### 2.3 Hydrogels with tunable mechanical dynamics

To rule out the effects of matrix stiffness on cell behaviors, we prepared PIC hydrogels with similar stiffness but with different mechanical dynamics by using different crosslinking strategies. Previous research of PIC models showed that intrafibrillar crosslinking didn’t significantly change the fibrous architecture, therefore not affecting gel stiffness significantly.[21, 33] We anticipate that the above three crosslinking strategies may mainly influence the mechanical dynamics of PIC hydrogels. To thoroughly investigate the mechanical properties of the hydrogels, rheological experiments, including temperature-induced gelation combined with SPAAC cross-linking reaction for PIC hydrogels, temperature-induced gelation for collagen, and ionic-crosslinking process for alginate, were performed *in situ* between the parallel plates of the rheometer.

For different PIC hydrogels, we employed a heating-cooling-reheating protocol[24] to compare their mechanical properties (Figure 2A). From the re-heating curve, PIC hydrogels with different crosslinking approaches showed similar shear storage modulus *G*′ at 37 °C (Figure 2B), which was in line with previous results.[21, 33] Specifically, *G*′ of pPIC, dPIC and cPIC was 370 ± 60 Pa, 320 ± 60 Pa and 290 ± 120 Pa, respectively (Figure S3B). However, the values of *G*′ at 5 °C were significantly different (Figure 2B, Figure S3A). PIC hydrogels with covalent crosslinking had the highest value (Figure 2B, solid rosy symbols), which is close to *G*′ at 37 °C, indicating that the thermoreversibility of highly crosslinked PIC hydrogels were entirely suppressed. However, PIC hydrogels with dynamic non-covalent crosslinking through catechol groups showed partially crosslinking (Figure 2B, purple solid symbols), and PIC hydrogels alone were completely thermoreversible (Figure 2B, blue solid symbols). Frequency sweep and strain sweep were tested to study the linear viscoelastic regime (Figure S4 and S5). To quantify the crosslinking difference, we calculate the ratio of the shear storage modulus at T = 5 and 37 °C as a phenomenological crosslinking efficiency. The crosslinking efficiency of PIC gels alone, PIC gels with dynamic non-covalent crosslinking, PIC gels with covalent crosslinking was 0.004, 0.0072, and 0.617, respectively (Table S1).

**Figure 2.**
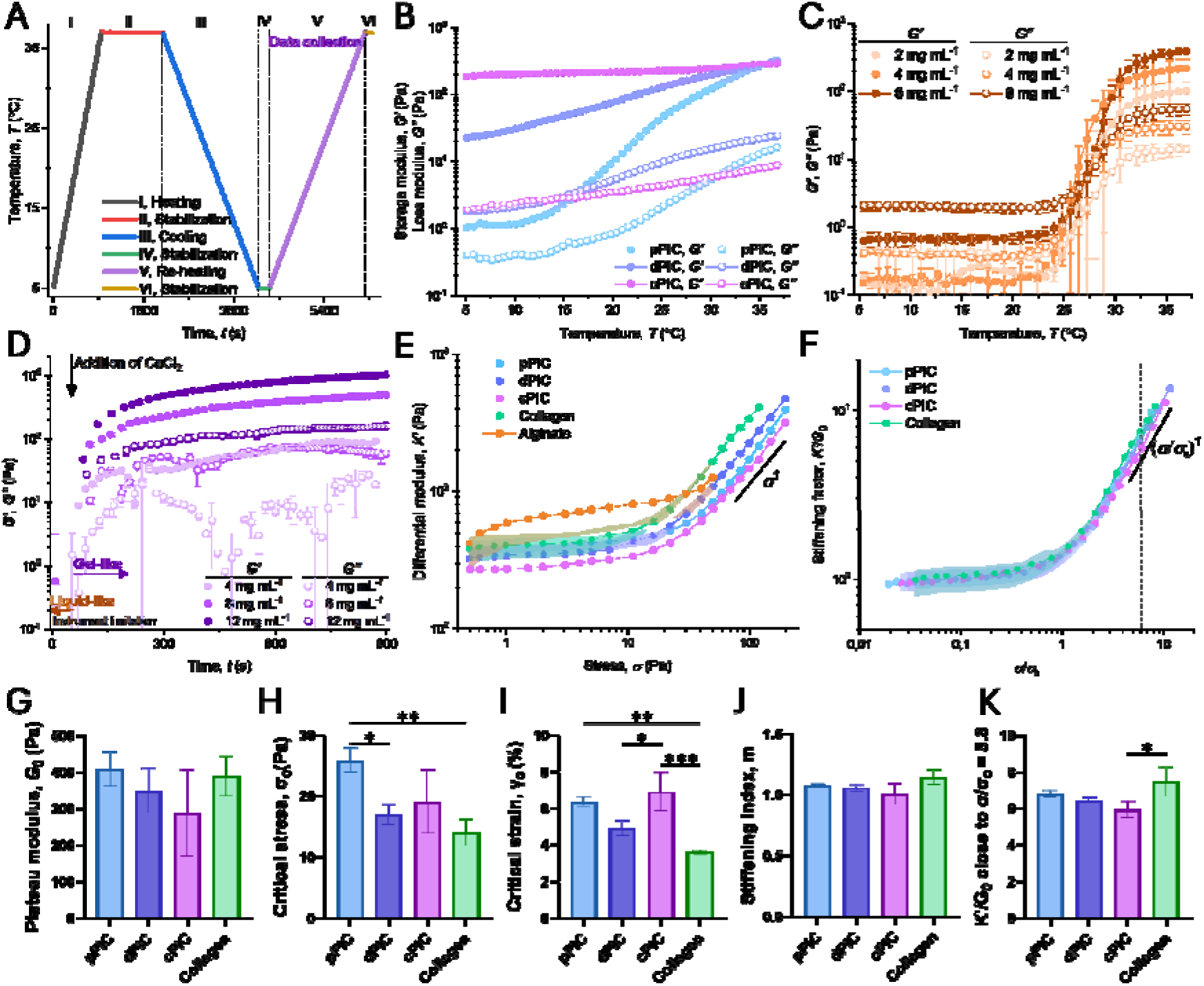
Mechanical properties of different hydrogels. (A) Crosslinking protocol. A freshly prepared solution of PIC with or without crosslinkers was placed on the rheometer at 5 °C, heated to 37 °C (stage I) and stabilized for 15 mins (stage II). Then the sample was cooled to 5 °C (stage III, cooling rate 1 °C min^−1^), shortly equilibrated (stage□), heated to 37 °C (stage, heating rate 1 °C min^−1^) and shortly equilibrated (stage□). (B) Linear viscoelastic regime: storage modulus *G*′ (solid symbols) and loss modulus *G’’* (open symbols) of different PIC hydrogels as a function of the temperature (blue: physically crosslinked PIC; purple: dynamically crosslinked PIC; rosy: covalently crosslinked PIC). Data are collected from stage. For all samples: concentration of PIC = 2 mg mL^−1^; the molar ratio of [linker]/[azide] is 0.4. (C) Storage modulus G′ (solid symbols) and loss modulus G’’ (open symbols) of collagen hydrogels as a function of the temperature. (D) Storage modulus *G*′ (solid symbols) and loss modulus *G’’* (open symbols) of alginate hydrogels as a function of the time under *in situ* ionic crosslinking. (E) Nonlinear mechanics: differential modulus *K*′ as a function of applied shear stress at *T* = 37 °C. (F) The modulus *K*′ versus the stress σ normalized with *G*_0_ and σ_c_, respectively, clearly showing similar stiffening response for the PIC hydrogels and collagen as a positive control. (G–K) Shear modulus *G*′ in the linear viscoelastic regime (G), critical stress σ_c_(H), critical strain γ_c_ = σ_c_/*G*′ (I), stiffening index *m* (J), and stiffening factor *K*′ /*G*_0_ at a normalized stress of 5.8 (K, data collection from the dash line in Figure 1F) of the PIC hydrogels and the collagen control gels. Error bars represent the standard deviation of the measurements for at least three times.

Moreover, the temperature-induced gelation for collagen hydrogels with different concentrations was investigated. With increasing concentrations of collagen, gels were stiffer ranging from 145 ± 30 Pa at 2 mg mL^−1^ to 430 ± 32 Pa at 6 mg mL^−1^ at 37 °C (Figure 2C, Figure S6). Additionally, the ionic crosslinking process of alginate hydrogels was monitored by the rheometer equipped with a home-made cup-shaped bottom plate,[21] wherein a calcium aqueous solution was added then diffused into the alginate polymer solution. After the addition of calcium solutions, alginate hydrogels were formed within a few minutes and gradually stabilized after 15 mins (Figure 2D). Strain sweep (Figure S7 and S9) and frequency sweep (Figure S8 and S10) were tested to study the linear viscoelastic regimes of collagen and alginate hydrogels, respectively. To prepare collagen and alginate hydrogels with similar *G*′ of PIC hydrogels at 37 °C, 6 mg mL^−1^ of collagen and 8 mg mL^−1^ of alginate hydrogels were used to evaluate their strain-stiffening properties.

Nonlinear strain-stiffening properties were investigated via the pre-stress protocol.[19] This approach involves applying a constant pre-stress σ combined with a small oscillatory stress to accurately capture the material’s response under load. The differential modulus *K*′ is determined from the ratio of the oscillatory stress to the resulting strain (K′=δσ/δγ). In the regime of low pre-stress, where the material behaves linearly, *K*′ named as plateau modulus *G*_0_ matches the conventional storage modulus *G*′. However, once the applied stress exceeds a critical threshold σ_c_, the system exhibits strong nonlinear stiffening, characterized by a sharp increase in modulus following *K*′∝σ^m^. Critical strain γ_c_ was determined by the ratio of σ_c_ and *G*_0_. The stiffening index *m* determines the intensity of the nonlinear stiffening response and is theoretically limited to values of *m* ≤ 3/2.

As expected, in the nonlinear regime, alginate hydrogels showed minimal stiffening response compared to PIC hydrogels and collagen (Figure 2E). Moreover, we found that intrafibrillar crosslinking didn’t significantly affect the stiffening response of PIC hydrogels, regardless of whether the crosslinking was dynamic non-covalent or covalent crosslinking. This effect became more evident when the data were normalized to *G*_0_ and σ_c_ (Figure 2F). Statistical analysis of most of the parameters, such as *G*_0_ (Figure 2G), σ_c_ (Figure 2H), γ_c_ (Figure 2I), and *m* (Figure 2J), showed no significant difference among different PIC hydrogels and collagen. Additionally, the stiffening factor *K*′/*G*_0_ determined at a set relative stress σ/σ_c_ = 5.8 showed no significant differences except covalently crosslinked PIC gels and collagen (Figure 2K). Therefore, the advantage of using fibrous hydrogels is to develop hydrogels with tunable mechanical dynamics by simply employing different crosslinking approaches without significantly changing the hydrogel concentration and stiffness.

### 2.4 Biocompatibility of the hydrogels

To assess the suitability of these hydrogel materials for *in vitro* cell culture, their biocompatibility was evaluated using hMSCs (Figure 3A). hMSCs were encapsulated in PIC hydrogels with different mechanical dynamics at a concentration of 2 mg mL^−1^, including pPIC, dPIC, and cPIC hydrogels (Figure 3B). Control conditions including strain-stiffening collagen and non-strain-stiffening alginate hydrogels were studied at a concentration of 6 mg mL^−1^ and 8 mg mL^−1^, respectively. To support cell– matrix adhesion, the PIC and alginate polymers used for cell encapsulation were functionalized with the GRGDS peptide (Gly–Arg–Gly–Asp–Ser), a widely used integrin-binding sequence, consistent with previous studies.[18, 34] PIC functionalization employed the same SPAAC chemistry used for cross-linking, utilizing additional azide groups on the PIC chains (Scheme S1). Alginate functionalization used carbodiimide crosslinker chemistry for GRGDS conjugation (details in methods). The biocompatibility of these hydrogels was confirmed using a multi-assay approach: a live–dead assay to assess cell viability, a viability assay based on metabolic activity, and a proliferation assay to quantify DNA.

**Figure 3.**
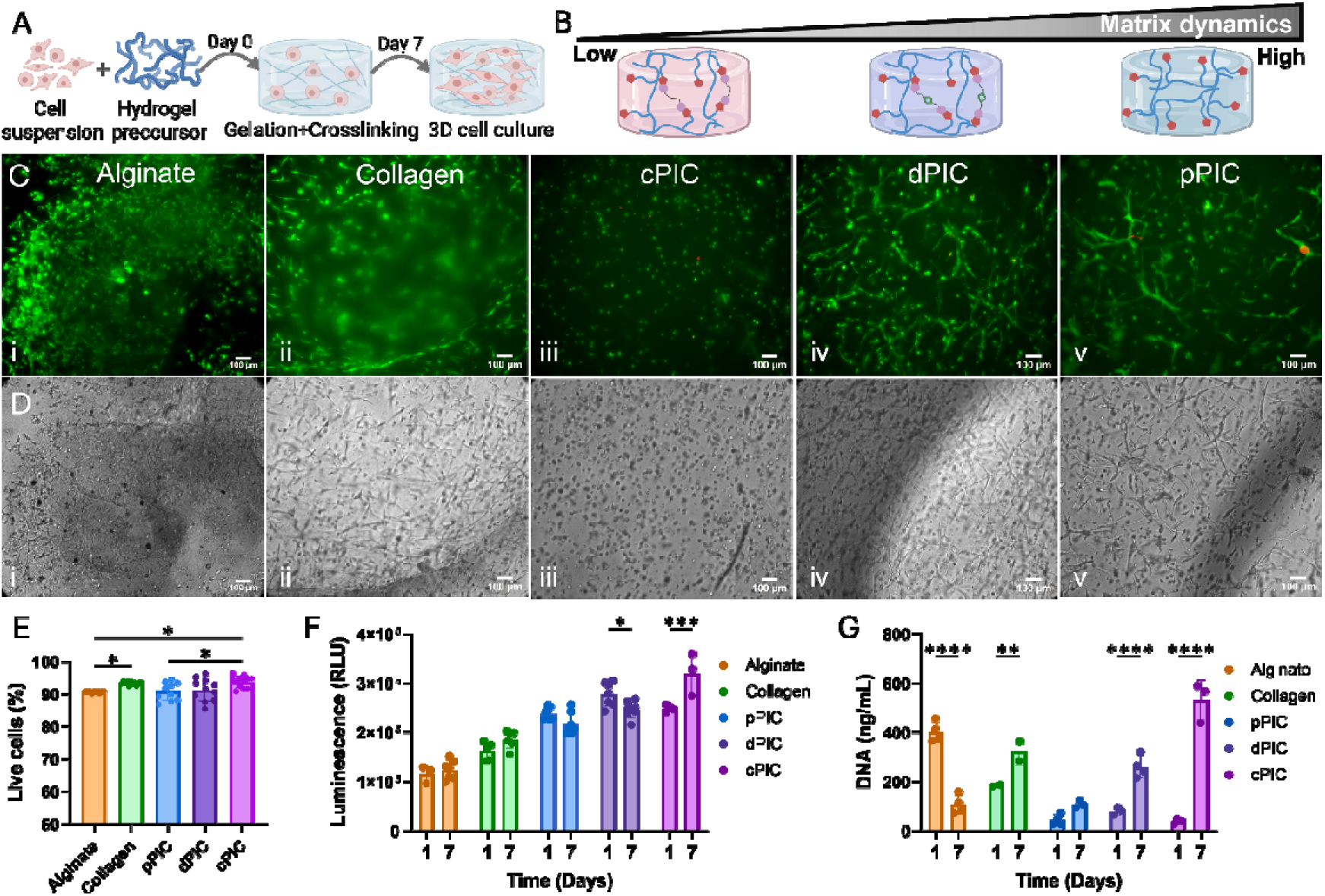
Biocompatibility of the hydrogels. (A) Scheme overview of the preparation of 3D *in vitro* models for cell culture. (B) Schematic illustration of PIC hydrogels with tunable mechanical dynamics by varying crosslinking strategies. (C, D) Fluorescent images (C) and bright field images (D) of hMSCs encapsulated in different hydrogels, including (i) alginate as a negative control, (ii) collagen as a positive control, (iii) covalently crosslinked PIC hydrogels, (iv) dual-dynamically crosslinked PIC hydrogels, and (□) physically crosslinked PIC hydrogels. Living cells (green) and dead cells (red) were stained with Calcein AM and EthD-1, respectively, at day 7. Scale bar = 100 μm. Conditions: cell density of hMSCs was 0.8 × 10^6^ cells mL^−1^, concentrations of alginate, collagen and PIC hydrogels were 8 mg mL^−1^, 6 mg mL^−1^, and 2 mg mL^−1^, respectively. For cPIC and dPIC hydrogels, the molar ratio of [linker]/[azide] were both 0.4. (E) Quantitative analysis of fluorescent images of live cells (green) and dead cells (red) after cell seeding at day 7. Data are presented as the percentage of live cells. (F) Metabolic activity of hMSCs encapsulated in different hydrogels at day 1 and 7. (G) DNA content in different hydrogels at day 1 and day 7. Statistics analysis was based on at least three samples. Data were tested through a one way or two-way ANOVA, significant differences (*, p < 0.05; **, p < 0.01; ***, p < 0.001; ****, p < 0.0001).

Live–dead staining results (Figure 3C) revealed high cell viability across all gel conditions. Green fluorescence from calcein-AM indicated live cells, while red fluorescence from thiazole red homodimer (TOTO-3) marked dead cells. In general, for all different hydrogel samples, cell viability was high at day 7, indicating that all tested hydrogel formulations supported robust cell survival. Quantitative image analysis showed viability ranging from 91% in alginate hydrogels ionically crosslinked with Ca^2^ ions to 94% in the covalently cross-linked cPIC gels (Figure 3E). Bright field imaging (Figure 3D) further supported these findings, displaying well-spread hMSCs except in covalently crosslinked cPIC hydrogels. It is important to note that PIC matrices are non-biodegradable and resist enzymatic degradation. Therefore, cell migration occurs via physical remodeling of the network, a mechanism previously observed in other non-degradable systems such as alginate hydrogels. The degree of crosslinking plays a critical role in modulating the process of physical remodeling, influencing the mechanical resistance and adaptability of the matrix to cells.[35] With increasing crosslinking degree, the matrix confinement was high, resulting in extremely limited cell migration.

The metabolic activities of hMSCs in different hydrogels was further investigated (Figure 3F). For alginate, collagen, and pPIC hydrogels, no significant differences were observed in the levels of metabolic activity between day 1 and day 7. Although metabolic activity reduction in dynamically crosslinked dPIC hydrogels on day 7 compared to that on day 1 was observed, its level was still higher than that in alginate, collagen and pPIC hydrogels. Interestingly, in covalently crosslinked cPIC hydrogels, metabolic activity significantly increased at day 7, indicating that cells in a less spread morphology may maintain higher proliferation. Besides, cell proliferation on day 1 and day 7 was quantified using a DNA assay, which measures DNA concentration as an indicator of cell numbers (Figure 3G). DNA content showed a substantial increase over time in these hydrogels except alginate. The observed reduction in DNA content in alginate hydrogels may be because the cell-laden gels are easily washed away during medium changes, leading to a loss of cells that is difficult to prevent. This issue could be addressed by integrating the hydrogels with cup-shaped electrospun scaffolds, which help retain the hydrogels and embedded cells. Notably, the covalently crosslinked cPIC hydrogels (Figure 3G, rosy columns) showed a dramatic increase in cell proliferation. Specifically, cPIC gels exhibited a 13.3-fold increase from day 1 to day 7, which was in line with the results of metabolic activities. Altogether these data reinforced the conclusion that all the hydrogel formulations support cell growth and are biocompatible.

### 2.5 Hydrogel mechanical dynamics regulates cell spreading

Matrix mechanical[29, 36] and dynamic properties[37] play an important role in regulating cell behaviors, thereby developing strategies to separate these two factors is necessary. By employing the intrafibrillar crosslinking approach in fibrous hydrogels with similar stiffness, we further investigated the role of matrix dynamics on cell behaviors, such as cell spreading. hMSCs with a cell density of 0.8 × 10^6^ cells mL^−1^ were encapsulated in different hydrogels. As matrix dynamics decreases with increasing crosslinking density, we hypothesize that the matrix network could modulate cell-matrix interactions. After cell encapsulation for 7 days, hMSCs were fixed and stained with phalloidin (F-actin) and DAPI (nucleus). We observed that hMSCs in pPIC gels (Figure 4A-, B-, C-, D-□) highly spread with the formation of cell-cell interaction network, while cells remained less stretched in dynamically non-covalently crosslinked dPIC gels (Figure 4A-, B-, C-, D-□), showing extended lengths and protrusions. In contrast, cellular morphology in covalently crosslinked cPIC hydrogels (Figure 4 A-, B-, C-, D-□) maintained spherical, indicating matrix dynamics could regulate cell spreading in a controlled manner. This phenomenon can be explained by the matrix remodeling capabilities in confined cellular microenvironments.[17] In such environments, cells navigate by squeezing through pores or creating channels by pulling and pushing on the fiber network. The degree of crosslinking plays a critical role in modulating this process, influencing the mechanical resistance and adaptability of the matrix to cells.[35] Additionally, cells cultured in collagen hydrogels (Figure 4 A-, B-, C-, D-□) showed more sprouting compared to those in dPIC and cPIC hydrogels, but didn’t form a cell network as observed in pPIC gels. Interestingly, cells in alginate hydrogels (Figure 4 A-, B-, C-, D-□) also formed a cell network, likely due to the folded gel structure caused during ionic crosslinking.

**Figure 4.**
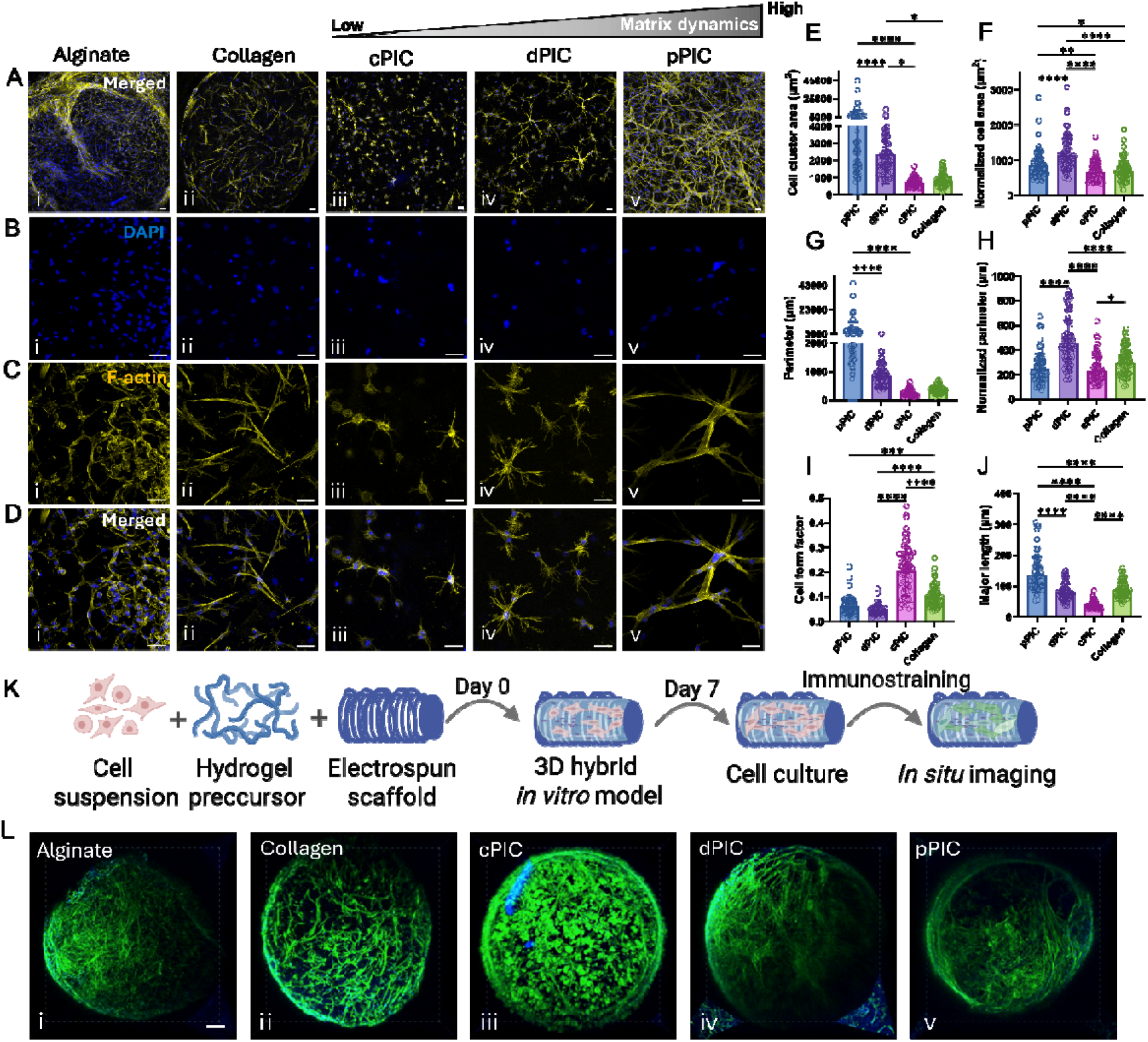
Matrix dynamics regulates hMSC morphology. (A-D) Immunofluorescent staining of hMSCs 3D cultured in different hydrogels for 7 days, including (i) alginate as a negative control, (ii) collagen as a positive control, (iii) covalently crosslinked PIC hydrogels, (iv) dual-dynamically crosslinked PIC hydrogels, and (□) physically crosslinked PIC hydrogels. Cells were fixed and stained with DAPI (blue) and phalloidin (yellow). Scale bar 100 μm. (E-J) Bar graphs about cell cluster area (E), normalized cell cluster area (F), cell perimeter (G), normalized cell perimeter (H), cell form factor (I), and major length (J). These parameters were quantified from the fluorescent images. Data were tested by one-way analysis of variance (ANOVA) followed by Tukey post hoc. *, p < 0.05; **, p < 0.01; ***, p < 0.001; ****, p < 0.0001; ns, not significant. Bars represent mean ± SD. (K) Schematic illustration of 3D hybrid *in vitro* models by integrating cell-laden hydrogels with electrospun scaffold. (L) Immunofluorescent staining of hMSCs 3D cultured in hybrid models for 7 days, including (i) alginate hydrocup, (ii) collagen hydrocup, (iii) cPIC hydrocup, (iv) dPIC hydrocup, and (□) pPIC hydrocup. Cells were fixed and stained with DAPI (blue) and phalloidin (green). Scale bar 200 μm. Conditions: cell density of hMSCs was 0.8 × 10^6^ cells mL^−1^. The concentrations of alginate, collagen and PIC hydrogels were 8 mg mL^−1^, 6 mg mL^−1^, and 2 mg mL^−1^, respectively. For cPIC and dPIC, the molar ratio of [linker]/[azide] were both 0.4.

Furthermore, to investigate cell morphological changes in response to matrix dynamics, we systematically quantified different key morphological features, including cell cluster area (Figure 4E), normalized cell area (Figure 4F), cell perimeter (Figure 4G), normalized cell perimeter (Figure 4H), cell form factor (analogous to circularity) (Figure 4I), major length (Figure 4J). Compared with cells cultured in other hydrogels, such as collagen (Figure 4D-ⅱ) and PIC-based hydrogels (Figure 4D-ⅲ, - ⅳ, and -ⅵ), cells encapsulated in alginate hydrogels exhibited more interconnected and irregular morphologies, with a tendency to form clustered structures (Figure 4D-i). Notably, no additional calcium ions were supplemented during culture, which may have contributed to alginate network destabilization and gel softening. Consequently, the alginate hydrogels developed folded and uneven structures during culture, compromising cell segmentation and morphological analysis and precluding meaningful quantitative comparisons among alginate, collagen, and the different PIC hydrogel conditions. Quantifications of (normalized) cell area showed hMSCs cultured in cPIC gels were morphologically similar to cells cultured in collagen, whereas they were significantly different from cells in pPIC and dPIC hydrogels (Figure 4E and 4F). Interestingly, although cell cluster area in pPIC gels was dramatically higher than in dPIC and cPIC gels (Figure 4E), the normalized cell area in these conditions was relatively close (Figure 4F). Similar phenomena were observed in these gel conditions for the (normalized) perimeter (Figure 4G and 4H). Cell form factor (Figure 4I) showed the highest level in cPIC gels, where cells were spherical. Surprisingly, despite cells spread more extensively in pPIC gels than in dPIC gels, there was no significant difference in their circularity. Additionally, the cellular major length (Figure 4J) in different PIC gels had similar results compared to cell cluster area and perimeter, where major length showed highest level in pPIC gels with high mechanical dynamics while lowest level in cPIC gels with low mechanical dynamics. Overall, by simply employing different crosslinking strategies, 3D *in vitro* PIC models with tunable mechanical dynamics without significantly changing stiffness may provide an advanced platform to investigate cell migration and spreading.

Next, we further studied cell spreading in hydrocups, where cell-laden hydrogels were injected into the hollow cup-shaped electrospun scaffolds (Figure 4K). An advanced fluorescent microscopy was used to evaluate cell spreading *in situ* instead of (cryo-)sectioning, allowing imaging cells without deformation with the depth up to a few millimeters. We observed that cells (green signals) in hydrogel matrices maintained similar morphology compared to those in gel alone, but spread more in the presence of electrospun scaffolds (Figure 4L, Supplementary Videos 1-5). It became clear that in hydrocups containing cPIC gels (Figure 4L-□), cells remain spherical in the center of hydrogels but spread more in the edge of electrospun scaffolds. It might be that the electrospun scaffolds were wetted by cell-laden polymer solutions during injection, allowing cell attachment on the electrospun scaffolds after hydrogel gelation. Electrospun scaffolds closely mimic the ECM with both microscale fiber size and porous architecture, which is beneficial for cells to receive these biophysical cues and in turn respond by increasing adhesion and spreading. Moreover, cells in the hydrocups containing alginate gels showed similar spreading morphology as other hydrocup conditions, further indicating that the introduction of electrospun scaffolds could help retain the gels and embedded cells.

### 2.6 Hydrogels as a toolbox for tuning cell secretome

Previous research demonstrated the onset of stress stiffening at critical stress σc levels as a key new indicator of stem cell differentiation, making a switch from adipogenesis to osteogenesis.[18] Notably, σc only showed a significantly difference between pPIC gels and dynamic non-covalently crosslinked dPIC gels (Figure 2H). To explore the potential of hydrocups as delivery systems for hMSCs as cytokine secretory factories, we investigated their cytokine release profiles. We chose different hydrogels to study the effects of their mechanical properties, such as strain-stiffening and mechanical dynamics on cell secretome, including strain-stiffening pPIC gels with high critical stress, dynamically crosslinked dPIC gels with low critical stress, collagen with low critical stress similar as dPIC gels as a positive control, and minimal strain-stiffening alginate as a negative control.

We quantified the secretion of 105 human cytokines from hMSCs cultured under the above four hydrocup conditions (Figure S11). The overall cytokine intensity distributions were broadly comparable among the four conditions, although pPIC and dPIC showed slightly higher average intensities than collagen and alginate (Figure 5A). However, hierarchical clustering and heatmap analysis revealed distinct cytokine secretion profiles across the four matrix conditions (Figure S12). The collagen condition displayed a cytokine pattern that was relatively distinct from the other three conditions, as indicated by the column-wise hierarchical clustering. In contrast, the alginate and pPIC conditions showed greater overall similarity and clustered more closely together, while dPIC exhibited an intermediate profile. Principal component analysis (PCA) further confirmed the separation among the conditions, with PC1 (47.5%) and PC2 (32.4%) explaining 79.9% of the total variance (Figure 5B). Collagen was primarily separated along PC1, whereas dPIC was distinguished along PC2, with pPIC and alginate positioned relatively closer together. Additionally, the cytokine Z-score distributions differed across the four matrix conditions (Figure S13). pPIC showed a clear shift toward positive Z-scores (Figure S13A), indicating relatively higher normalized levels for many cytokines, whereas collagen was predominantly shifted toward negative Z-scores (Figure S13C), indicating relatively lower levels. In comparison, dPIC (Figure S13B) and alginate (Figure S13D) exhibited broader and more balanced distributions across negative and positive Z-scores. Together, these complementary analyses demonstrate that the matrix properties influence the overall cytokine secretion profile.

**Figure 5.**
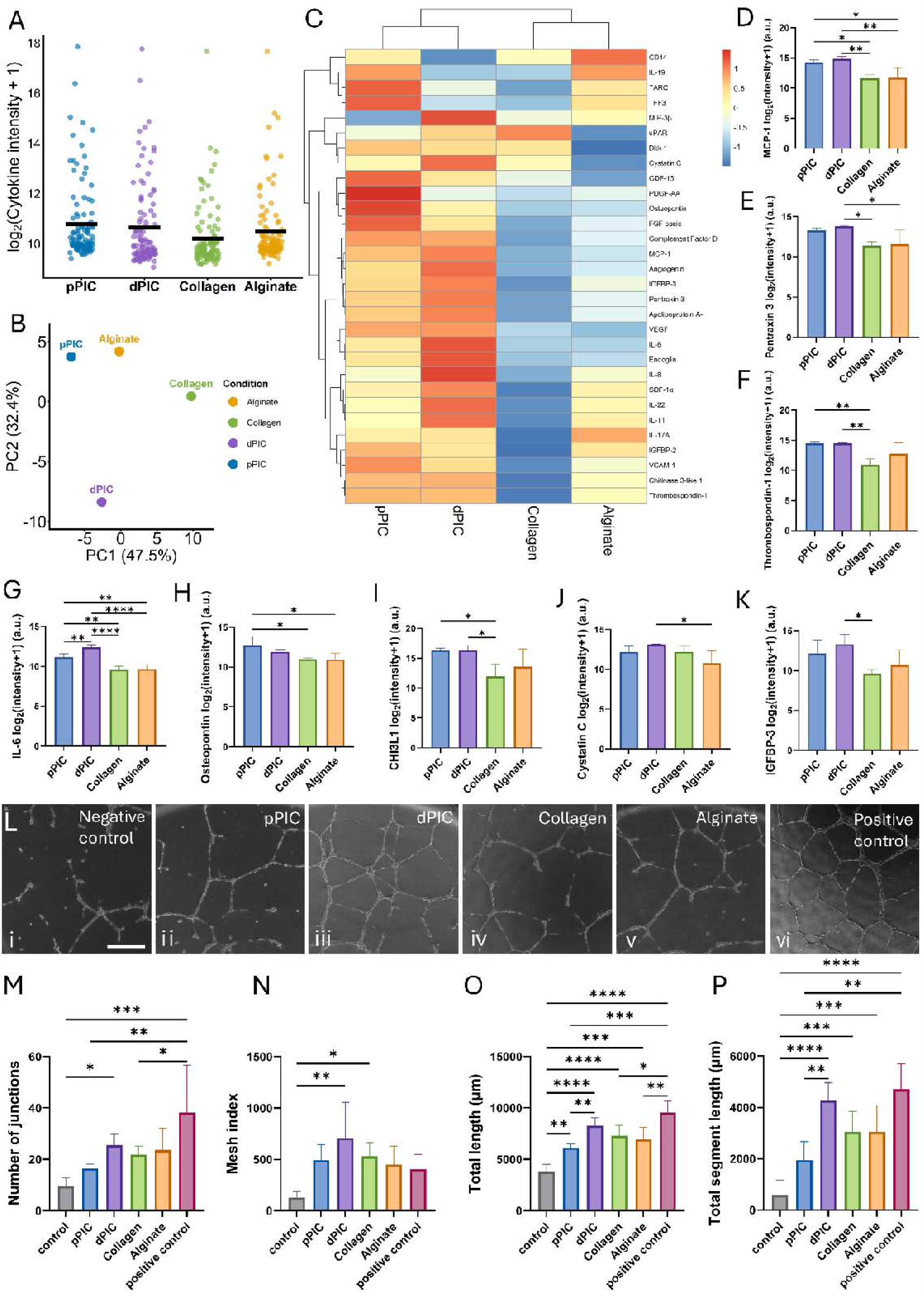
Cytokine secretion of hMSCs cultured in hydrocups. (A) Dotplot visualization of cytokine intensity levels across the conditions. Black lines represent the average level. (B) PCA of cytokine profiles across different matrix conditions, based on the full cytokine intensity profiles of pPIC, dPIC, collagen, and alginate conditions. (C) Clustered heatmap showing hierarchical clustering of the 30 cytokines exhibiting the greatest variation across the four matrix conditions. Cytokine levels were row-scaled to visualize relative differences among collagen, dPIC, pPPIC, and alginate conditions, , according to the chemiluminescence scanner images of cytokine secretion analysis shown in Figure S11. An overview of the clustered heatmap of the 105 cytokines is shown in Figure S12. Red and blue colors indicate relatively higher and lower levels, respectively, for each cytokine. The displayed Z-score color scale is approximately −1.5 to +1.5. Four data sets were averaged. (D-K) Intensity of the expressed angiogenesis-releated cytokines, including (D) MCP-1, (E) Pentraxin 3, (F)Thrombospondin 1, (G), (H), (I), (J), and (K). (L) HMVEC tube formation assay under six media treatments, including hMSC basic proliferation culture medium as a negative control (i), conditioned media from hMSC in pPIC hydrocup (ii), dPIC hydrocup (iii), collagen hydrocup (iv), alginate hydrocup (□), and HMVEC proliferation medium as a positive control (□). Scale bar 100 μm. Note, in the first five conditions, 50% of HMVEC proliferation media were supplemented. (M-P) Quantification analysis of the tube formation assay to compare the differences in the tube-forming ability of HMVEC. Parameters of numbers of junctions (M), mesh index (N), total tube length (O), and total segment length (P) were analyzed from images in panel G. Statistics analysis was based on at least three samples. Data were tested through a one-way ANOVA, significant differences (*, p < 0.05; **, p < 0.01; ***, p < 0.001; ****, p < 0.0001).

We further analyzed the hierarchical clustering of the 30 most variable cytokines, demonstrating distinct condition-dependent secretion profiles (Figure 5C). The column dendrogram separated the four matrix conditions into two major clusters, with dPIC clustering with pPIC and collagen clustering with alginate, indicating greater similarity in their overall cytokine profiles within these pairs. At the individual cytokine level, condition-dependent patterns were evident, with subsets of cytokines showing relatively increased or decreased levels under specific matrix conditions. In particular, pPIC showed relatively elevated levels across a substantial cluster of cytokines, whereas dPIC exhibited increased levels in a different subset.

Of interest, the expressed angiogenesis-releated proteins,[38–40] like vascular endothelial growth factor (VEGF), monocyte chemoattractant protein 1 (MCP-1), interleukin-8 (IL-8), thrombospondin 1, and pentraxin 3 were identified. Pro-angiogenic factor like MCP-1[41] exhibited a significant increased expression in PIC-related hydrocup conditions (Figure 5D), but there was no significant difference between the conditions of pPIC and dPIC hydrocups. Moreover, pentraxin 3 mediates angiogenesis, which can be pro-angiogenic[42] or anti-angiogenic[43], depending on its molecular interactions and local microenvironment. Interestingly, pentraxin 3 was significantly expressed in dPIC hydrocup conditions (Figure 5E). Although there was no significant difference between pPIC and dPIC hydrocup conditions, MCP-1 and pentraxin 3 were expressed more in dPIC hydrocups. Anti-angiogenic factor such as thrombospondin-1[40] exhibited a decreased expression in collagen but an increase in other three conditions (Figure 5F). Although no significant difference was observed across other angiogenesis-related factors, the PIC-based matrices tended to exhibit relatively higher levels of several factors associated with endothelial activation, migration, and vascularization (Figure S14).[44–46]

In addition to angiogenesis-related cytokines, differences were also observed in other cytokine categories, including pro-/anti-inflammatory,[38, 47, 48] as well as cytokines involved in cell migration[49], proliferation[50], and tissue remodeling.[51] Interleukin-6 (IL-6, Figure 5G), a proinflammatory cytokine, and osteopontin (Figure 5H) which induce angiogenesis[52, 53] exhibited relatively higher levels in the PIC-based conditions, similar as GDF-15 expression, a pleotrophic cytokine with roles in inflammation, cancer and metabolism[54] (Figure S15). Previous results demonstrated that the expressed pro-inflammatory chitinase-3-like 1 (CHI3L1) cytokines[55] were secreted significantly more in the alginate hydrocups compared to the 2D cell culture.[16] Interestingly, CHI3L1 cytokines were significantly secreted at high levels in PIC-related hydrocups compared to collagen and alginate hydrocups (Figure 5I). Cystatin C, a cysteine-protease inhibitor that regulates cathepsin activity and thereby contributes to proteolytic activity and ECM turnover,[51] was relatively comparable among pPIC, dPIC, and collagen but lower in alginate (Figure 5J). IGFBP-3, a regulator of IGF signaling involved in the modulation of cell proliferation, survival, and vascular responses,[56] was relatively higher in dPIC and lower in collagen (Figure 5K). IL-17A and IL-22 cytokines contributing to cell migration[49] were downregulated in collagen hydrocups (Figure S16). Complement factor D cytokines, which have been demonstrated as a strategic target for advancing therapeutic control of pathological complement activation,[57] were expressed more in PIC-related hydrocups (Figure S17).

Overall, we demonstrated that cell cytokine secretion can be modulated by selecting different hydrogel formulations and further tuning their mechanical properties. Different hydrogel types established distinct physicochemical and mechanical microenvironments, which were associated with differences in matrix stress-stiffening behavior and mechanical dynamics. Importantly, modulation of these mechanical characteristics within and across hydrogel formulations further influenced cytokine secretion, indicating that the cellular secretory response is governed not only by hydrogel composition but also by the gel mechanical dynamics. These findings highlight hydrogel selection as an important determinant of the cellular microenvironment, while demonstrating that mechanical properties provide an additional and tunable tool to regulate cell secretion. Collectively, our work provides a versatile strategy for engineering hydrogel microenvironments to modulate cytokine secretion.

### 2.7 Tube formation assay for evaluating the functionalities of cell secretome

To assess the functionalities of the cytokines secreted by hMSCs, we performed a tube formation assay by comparing different conditioned media collected from hMSC culture. hMSC and human microvascular endothelial cells (HMVEC) proliferation culture media were used as negative and positive controls, respectively. The HMVEC tube formation ability in conditioned medium collected from dPIC hydrocups (Figure 5L-) was dramatically enhanced comparing to other conditioned groups, including pPIC (Figure 5L-□), collagen (Figure 5L-□), and alginate hydrocups (Figure 5L-□), showing the formation of well-connected capillary-like structures, which was in line with the results of angiogenesis-related cytokine analysis. The simultaneous detection of both pro- and anti-angiogenic factors in the cytokine array may indicate that hMSC-mediated regulation of angiogenesis is highly dependent on cellular local microenvironment, rather than fitting neatly into a strictly pro- or anti-angiogenic classification. In the negative control group, the HMVEC tube formation ability was significantly attenuated (Figure 5L-□).

Furthermore, we systematically quantified four key features, including numbers of junctions indicating complexity and connectivity of the vascular-like network (Figure 5M), mesh index reflecting stability and continuity of the network (Figure 5N), total tube length representing overall outgrowth capability (Figure 5O), and total segment length giving insight into branching complexity and fine extension (Figure 5P). The quantification analysis showed that there was no significant difference in tube formation ability between cells cultured in dPIC-treated medium (Figure 5M-P, purple columns) and endothelial cell proliferation medium as a positive control (Figure 5M-P, red columns). These findings illustrate that the hMSC secretome remains functionally active, and the secreted cytokines can promote HMVEC tube formation without impairing the cytokine-mediated signaling, leading to angiogenesis.

In summary, we developed a versatile hydrogel-based platform that enables the regulation of hMSC secretory behavior through the simple and tunable modulation of hydrogel mechanical dynamics but with similar stiffness at their initial gel status. By independently controlling the matrix mechanical properties, this platform allows the mechanical microenvironment experienced by hMSCs to be tailored, thereby providing a strategy to regulate their secretome. This mechanical regulation offers a facile strategy for engineering the cellular microenvironment without requiring changes in cell source or exogenous biochemical stimulation. More broadly, the ability to modulate hMSC-derived paracrine signaling through tuning matrix mechanics may provide a generalizable approach for optimizing cell-based therapeutic functions and advancing the development of next-generation regenerative strategies.

## 3. Conclusion

We have prepared a hydrogel library composed of synthetic, highly biomimetic fibrous PIC hydrogels with tunable mechanical dynamics by simply employing different crosslinking approaches, including physically, dynamically dual, and covalently crosslinking, respectively. Unlike other materials where the mechanical properties, particularly the stiffness, typically are significantly changed by the addition of crosslinkers, in these materials the stiffness is similar due to the intrafibrillar crosslinking. As a result, the intrafibrillar crosslinked hydrogels show similar stiffness but tunable mechanical dynamics without significantly changing the hydrogel components, exhibiting a direct correlation between cell morphology and matrix mechanical dynamics. Moreover, a 3D hybrid *in vitro* model is developed with combination of cup-shaped hollow electrospun scaffold and hydrogels to mimic cell-surrounding fibrous matrix.

We then demonstrate that the cytokine secretion profile of hMSC can be regulated through simply changing the hydrogel types and/or mechanical dynamics, leading to a tunable tube formation ability of endothelial cells cultured with the conditioned media from hMSCs. It is imperative to note that the degree of tube formation is contingent on the type of conditioned culture media. However, due to the hydrogel contraction during cell culture, a relatively low cell density was employed in our work to maintain the structural integrity of the constructs. This relatively low cellularity may have contributed to the limited levels of cytokines detected in the conditioned media, making it challenging to identify potential comprehensive differences in cytokine secretion among the different gel conditions. Future studies could therefore explore increasing the cell density within the gels as an initial approach to enhance the overall production and accumulation of secreted cytokines, thereby enabling more sensitive characterization of the cytokine secretion profiles. Despite this limitation, our findings underscore the significance of hydrogel properties in modulating cell behaviors, such as morphology and secretome production. We anticipate that our findings can be adapted to other fibrous hydrogels, either natural or synthetic, which may offer a promising approach for advancing regenerative strategies via regulating cell-matrix interactions.

## 4. Materials and Methods

### 4.1 Electrospun scaffold preparation

Cup-shaped electrospun scaffolds were produced using 300PEOT55PBT45 (PolyVation), synthesized with 300 kDa poly(ethylene glycol), with a 55/45 (w/w) PEOT/PBT ratio. A 25% (w/v) solution of 300PEOT55PBT45 was prepared by overnight dissolution in 30% (v/v) 1,1,1,3,3,3-Hexafluoro-2-propanol AR (HFIP) (Bio-Solve) and 70% (v/v) Chloroform (Sigma-Aldrich) under agitation at room temperature. The polymer solution was electrospun on a 1.8 mm diameter mandrel rotating at 100 RPM. The mandrel was attached on the motor on one side and open on the other side. The open side had a rounded tip to prevent charge accumulation at focal points. The polymer solution was fed to a charged needle at 3 mL/h in a chamber of 23-25 °C and 30-40% relative humidity. The needle, charged between 10-15kV, was positioned at a roughly 30° angle to increase coverage on the open end of the mandrel. The mandrel was charged -1.5kV. After an initial spinning period of ∼1min in the middle of the mandrel, the needle was moved to the side to spin on the open end of the mandrel. The working distance was around 18 cm, depending on the final position of the needle. The electrospun scaffolds were produced in 2 min. The hydrocups were removed from the mandrel with 70% (v/v) ethanol, dried and sterilized for 2 hours with 254 nm UV light in vacuum.

### 4.2 Hydrogel preparation

The synthesis of PIC polymers was performed with a minor modification.[20] Briefly, azide-functionalized PIC was obtained by copolymerization of a methoxide and an azide-appended monomer (ratio OMe to N_3_ is 87:13) in toluene in the presence of Ni (ClO_4_)_2_·6H_2_O solution as catalyst. A monomer:catalyst ratio of 1:1000 was used to prepare short PIC polymers. The polymerization mixture was stirred overnight, and full monomer conversion was confirmed by the disappearance of the characteristic isocyanide peak at 2140 cm^−1^ in the infrared spectrum. The polymers were isolated by precipitation in diisopropyl ether for three times and air drying, after which they were stored at −20 °C until use. For biofunctionalization with the cell adhesive peptide GRGDS peptide (Gly–Arg–Gly–Asp–Ser), an earlier published protocol was used,[21, 23] where the peptide is equipped with a dibenzocyclooctyne (DBCO)-terminated spacer and subsequently clicked to the PIC polymers using the highly efficient strain-promoted azide-alkyne cycloaddition (SPAAC) reaction.

PIC pellets were dissolved at a concentration of 4 mg mL^−1^ in phosphate-buffered saline (PBS) as a stock solution at 4 °C overnight. Physically crosslinked PIC hydrogels were prepared by heating up an aqueous cold PIC polymer solution above its gelation temperature *T*_gel_, resulting in PIC chain aggregation into bundles, thus forming a thermo-responsive hydrogel network through hydrogen bonding. Dual physically crosslinked PIC hydrogels were prepared in a similar way as single physically crosslinked PIC hydrogels, but by addition of catechol-DBCO crosslinkers into the PIC polymer solution. After mixing, the solution was heated to promote the SPAAC cross-linking reaction between PIC hydrogels and the crosslinkers, leading to the formation of catechol-functionalized PIC hydrogels. To prepare chemically crosslinked PIC hydrogels, DBCO-PEG4-DBCO crosslinkers were added to an aqueous cold PIC polymer solution, followed by heating. PBS buffer was used to dilute the PIC stock solution to desired concentrations. Collagen type hydrogels with high protein were prepared following the preparation protocol from Corning. Briefly, a neutralization buffer solution of 10× PBS, 18 mM NaOH, and miliQ water was used to neutralize the pH of collagen stock solution. Collagen hydrogels were formed under gelation at 37 °C for half an hour.

Alginate was purified using a previously published protocol. Briefly, 1% (w/v) alginate was dissolved overnight under agitation in ultrapure water (18 MΩ, Milli-Q UltraPure Water System, Millipore). The solution was filtered through a series of different diameter membrane pores (1.2, 0.45 and 0.22 μm, VWR), freeze-dried and stored at −20 °C. To allow cell adhesion to the alginate, it was modified with GRGDS peptide by using aqueous carbodiimide chemistry (EDC chemistry). Briefly, the purified alginate was dissolved at 1% (w/v) in 0.1 M 2-(N-morpholino) ethanesulfonic acid (MES) buffer solution (0.1 M MES buffering salt, 0.3 M NaCl, pH adjusted to 6.5 using 1 M NaOH, Sigma). N-Hydroxy-sulfosuccinimide (sulfo-NHS, Pierce Chemical) and 1-ethyl-(dimethylaminopropyl)-carbodiimide (EDC, Sigma, 27.40 mg per g alginate), at a molar ratio of 1:2, was then added to the solutions, followed by 65.9 μmol RGD for every gram alginate. This mixture was incubated under agitation for 20h and subsequently quenched with 18 mg of hydroxylamine hydrochloride (Sigma) per gram alginate. The final product was dialyzed over three days at 4 °C (MWCO 3500, Spectra/Por®, VWR) against decreasing concentrations of NaCl in ultrapure water, freeze-dried and stored at -20 °C. The RGD modified alginate is referred to as “alginate” throughout the manuscript.

### 4.3 Rheology measurements

The mechanical properties were analyzed on a Discovery HR-2 rheometer (TA Instruments) with a geometry gap of 500 µm. A pre-cold PIC polymer solution at a concentration of 2 mg mL^−1^ and collagen polymer solutions were loaded onto the rheometer at 5 °C, respectively, followed by heating up to 37 °C and stabilization. Once the hydrogel network was formed, the temperature was lowered to 5 °C. At this point, non-crosslinked PIC reverts to a soluble solution, while the crosslinked samples (partially) retained their gel structures. Subsequently, all samples were re-heated to 37 °C. For long time strain-stiffening experiments, a layer of silica oil was added to seal the gap to prevent sample dehydration. Then the gel solutions were heated with a controlled rate (2 °C min^−1^) to the desired temperature. The mechanical properties in the linear viscoelastic regime (strain amplitude γ = 4 %, frequency ω = 1 Hz) were recorded, unless noted otherwise. The nonlinear response of the hydrogels was evaluated at a set pre-stress σ0 (with σ < 0.1 σ0) with a small stress amplitude by applying frequency sweep (0.1–10 Hz). To monitor the crosslinking process of ionically crosslinked alginate hydrogels, we used a home-built cup-shaped bottom plate, wherein a calcium ion solution at a desired concentration can be placed after the geometry is closed, which then diffuses into the alginate polymer network, resulting in ionic crosslinking. Each sample was measured at least three times to ensure repeatability.

### 4.4 Cell culture and encapsulation

hMSCs were from a woman donor of 30 years old (PromoCell). hMSC at passage 2 were expanded at cultured in MEM Alpha Medium with GlutaMAX (32561-029, Gibco), supplemented with 10% (v/v) of fetal bovine serum (FBS), 1% penicillin/streptomycin solution (Pen-Strep) and 1% ascorbic acid-2-phosphate until 80% confluency was reached. Cells passaged at 5 were used for cell culture experiments. For the seeding process, cells were collected using 0.05% trypsin-EDTA treatment, then resuspended in the medium after centrifugation. The RGD-conjugated PIC polymers were dissolved in medium at 4 °C for overnight, and the polymer solution at a concentration of 2 mg mL^−1^ was sterilized by using 20 µm of membrane filter. Cells with a final density of 0.8 × 10^6^ cells mL^−1^ were mixed with the above polymer solution and seeded in a 15-well microplate (uncoated, Ibidi GmbH, Martinsried, Germany) with a volume of 10 µL each well to study the biocompatibility of CPIC hydrogels and the hMSCs spreading in hydrogels with different mechanical dynamics. After seeding, cells were 3D encapsulated in the hydrogels during incubation at 37 °C, 5% CO_2_ for 15 min. Medium of 50 µL was added on top of the hydrogels, and was refreshed every two or three days. As a control, cells were seeded in alginate and collagen hydrogels.

### 4.5 Live-dead staining

To analyze cell viability in the hydrogels, a live-dead assay was carried out at day 7. A staining solution of 2.5 μm of ethidium homodimer-1 (EthD-1, E1169, Thermo Fisher) and 1 μm of calcein acetoxymethyl (Calcein AM, c3099, Thermo Fisher) was prepared in medium, followed by adding to cells encapsulated in hydrogels. Hydrogels were washed 3 times with αMEM without phenol red (Gibco), immersed in PBS, and imaged directly with a fluorescence microscope (Nikon Ti-E microscope, Japan).

### 4.6 Metabolic activity

Metabolic activity of hMSCs was quantified using the Cell Titer-Glo 3D assay (Promega) on day 1, and 7. Specifically, equal volumes of reagent and medium were added into the hMSCs laden hydrogels then mixing them by pipetting. After 30 minutes incubation at room temperature, 100 µL of supernatant was transferred to a white bottom 96 well plate and luminescence was measured by a CLARIO star plate reader (BMG Labtech, Germany).

### 4.7 DNA quantification

Cell extraction from hMSCs laden hydrogels were obtained by freeze-thaw 3 times, digested overnight at 56 with 1 mg mL^−1^ proteinase K and freeze-thaw 3 more times. DNA quantification was performed following the protocol provided by the CyQuant cell proliferation assay kit (Thermo Fisher, USA). Briefly, the supernatant from above cell extraction was further incubated with lysis buffer containing RNase for 2 hours at room temperature. An equal volume of 2× GR-dye solution was added to mixture followed by incubation for 15 minutes. Samples (200 μL) were transferred to a black 96-well plate and the fluorescence at 520 nm were measured by a CLARIO star plate reader.

### 4.8 Immunofluorescence staining

For immunofluorescence analysis, encapsulated cells in hydrogels or hydrocups at day 7 were fixed in 4% paraformaldehyde in pre-warmed PBS for 10 min at room temperature, followed by pre-warmed PBS washing for 3 times. Cells were permeabilized and blocked with 1% (v/v) Triton X-100, 0.05% (v/v) tween-20, 5% (v/v) goat serum, and 1% w/v bovine serum albumin (BSA) in PBS for 2 h. Phalloidin (Alexa Fluor 647, ThermoFisher Scientific, Cat #10656353) and 4′,6-diamidino-2-phenylindole (DAPI, ≥ 95.0%, Sigma–Aldrich, Cat #32670) was diluted to 1:100 and 1:200 in 1% w/v bovine serum albumin (BSA, WVR, Cat #421501J) in PBS, respectively. After removing the blocking buffer, phalloidin was added to the cells for 60 min at room temperature, followed by PBS washing for three times. Then cells were incubated with DAPI for 20 min at room temperature, and rinsed with PBS for three times. For cells encapsulated in hydrogels and hydrocups, cells were imaged by using a TCS SP8 STED confocal microscope and a confocal microscope (Nikon Eclipse Ti2 2-photon), respectively.

### 4.9 Morphological analysis

Immunofluorescence images were analyzed based on the actin and nuclei channels using ImageJ. Several shape-dependent cellular and nuclear morphological features were assessed, including cell cluster area, normalized cell area, cell perimeter, normalized cell perimeter, cell form factor (analogous to circularity), and major length.[58]

### 4.10 Cell secretome

To measure cytokine secretion from hMSCs, cells were cultured in cell-laden hydrocups for 6 days after seeding. After this, medium was refreshed with exactly 1 ml basic medium and incubated with the cells for 24 hours. In this medium, 105 human cytokines were measured simultaneously using the proteome profiler human XL cytokine array kit (R&D systems) according to the manufacturer’s protocols. Briefly, the cytokine containing medium was incubated on PVDF membranes containing spots with antibodies for each cytokine. A mix of primary antibodies was incubated, followed by horseradish-peroxidase -coupled secondary antibody incubation. Images were captured at different timings in order to optimize the signal-to-background ratio of the image by using the chemiluminescence scanner Azure Biosystems C600. All membranes were imaged at the same time to allow relative comparison between conditions.

Data were analyzed through ImageJ to read the cytokine intensity from the membranes. PCA of cytokine profiles, hierarchical cluster analysis with complete clustering method, and dotplot visualization of cytokine intensity levels were performed with R software version 4.4.0. Regarding the clustered heatmap, log_2_-transformed cytokine intensities were standardized to Z-scores across conditions for each cytokine. Hierarchical clustering was performed for both experimental conditions (columns) and cytokines (rows) using Euclidean distance and Ward’s D2 linkage method, and the resulting dendrograms were displayed along the corresponding dimensions of the heatmap. Statistical analysis was performed with Graphpad Prism version 9.3.1.

### 4.11 Tube formation assay

Geltrex (LDEV-Free Reduced Growth Factor Basement Membrane Matrix, Gibco) was used to coat the 15-well ibidi slide (10 µl per well) and was incubated at 37 °C for 30 min. Next, HMVECs were seeded on top at a density of 4 × 10^4^ cells cm^−2^ in EGM-MV2, followed by culturing of overnight under different conditioned media collected from hMSC culture, hMSC and HMVEC proliferation culture media, respectively. After 16 h, the tube formation network was imaged with brightfield using a fluorescence microscope (Nikon Eclipse Ti-E).

For each well, three brightfield images were taken for analysis. Fiji (angiogenesis analyzer plugin) was used to assess vascular networks, which detect extremities (extremities in binary skeleton), junctions (joined nodes, pixels with at least 3 neighbors), branches (elements delimited by a junction and one extremity), segments (elements delimited by two junctions), and meshes (areas enclosed by segments).[59] Next, analyze the number of junctions, mesh index, total length, and total segment length in each image. For each group, the averaged values from all images were used to represent its 2D tube formation results.

### 4.12 Statistical Analysis

Statistical analyses of the values for all experiments were presented as mean ± standard deviation. Statistical analysis was based on at least three samples using one-way analysis of variance (ANOVA), followed by the Tukey’s multiple comparison tests. Probability values p > 0.05 were considered not significant. For significant differences: * p < 0.05, ** p < 0.01, *** p < 0.001, **** p < 0.0001.

## CRediT authorship contribution statement

**Wen Chen:** Writing – review & editing, Writing – original draft, Visualization, Methodology, Investigation, Formal analysis. **Hao Wu:** Writing – review & editing, Methodology, Investigation. **Zhengdong Gao:** Writing – review & editing, Methodology, Investigation. **Hongjuan Weng:** Writing – review & editing, Methodology, Investigation. **Behrad Shaghaghi:** Writing – review & editing, Methodology, Investigation. **Katrien V. Bernaerts:** Writing – review & editing, Resources, Funding acquisition. **Paul H. J. Kouwer:** Writing – review & editing, Resources. **Paul Wieringa:** Writing – review & editing, Resources, Funding acquisition. **Lorenzo Moroni:** Writing – review & editing, Supervision, Resources, Project administration, Funding acquisition, Conceptualization.

## Acknowledgements

This work was supported by the European Research Council (ERC, no. 101057070); the Dutch Research Council (NWO) grant (no. TTW 18647 – VICI); and the China Scholarship Council (CSC, no. 202206170025, no. 202206220044, and no. 202006220058). All illustrations were created by BioRender.com.

