## Supplementary Information for "Hydrogel chemistry modulates mesenchymal stromal cell outgrowth and cytokine secretion profile"

**Additional Figures**

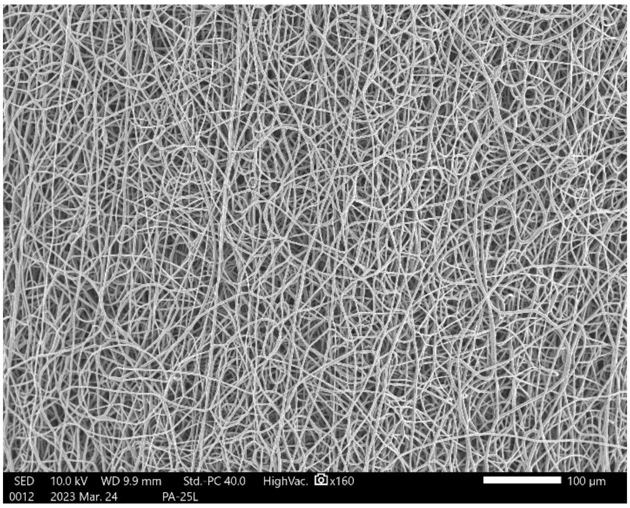


**Figure S1.** SEM image of the electrospun scaffold. Scale bar 100 µm.

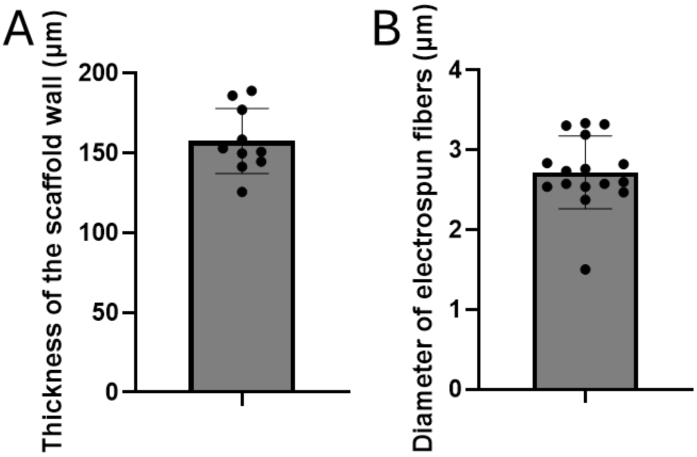


**Figure S2.** Quantification analysis of the electrospun scaffolds. (A) The thickness of the electrospun scaffold wall. (B) The diameter of the electrospun fibers.

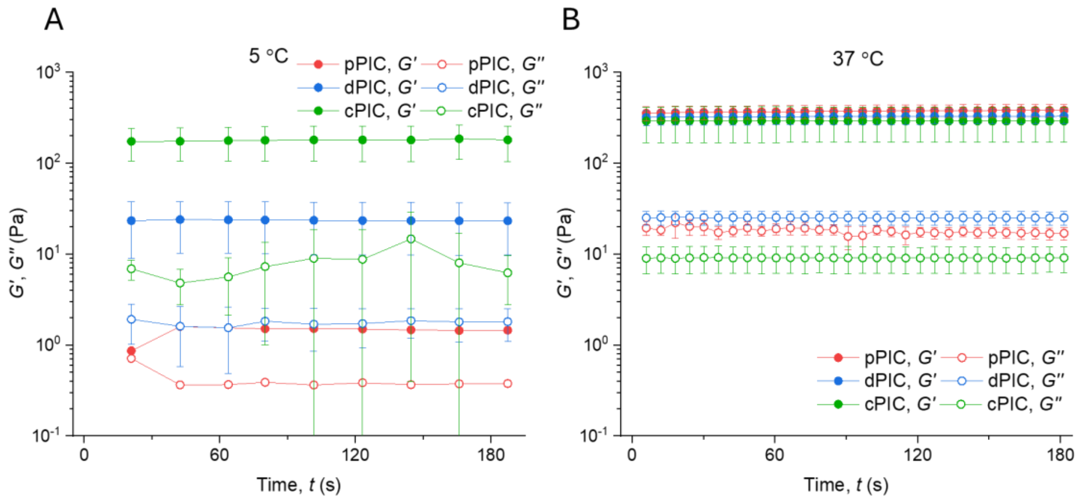


**Figure S3.** Time sweep of PIC hydrogels with different crosslinking approaches at 5 °C (A) and 37 °C (B). Conditions: For all samples: concentration of PIC = 2 mg mL$^{-1}$; the molar ratio of [linker]/[azide] is 0.4.

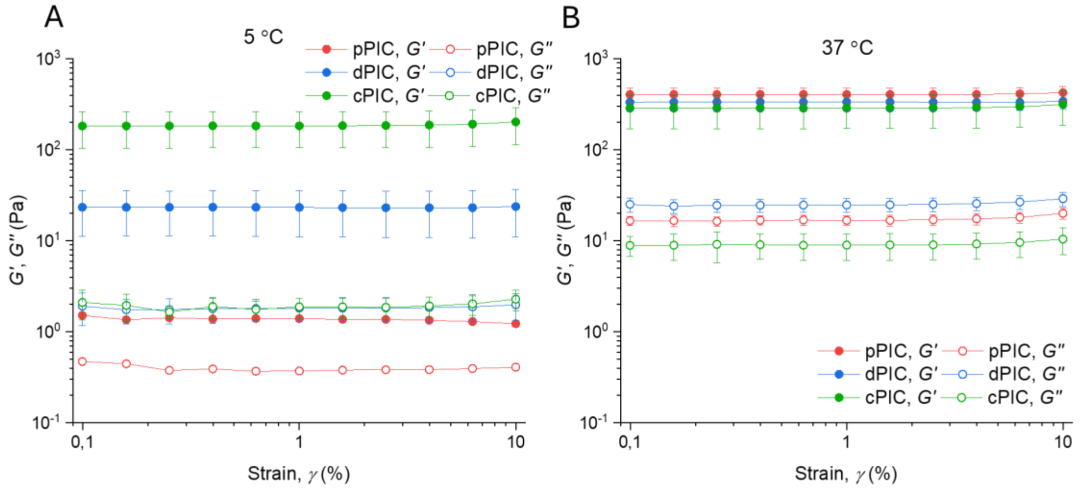


**Figure S4.** Viscoelastic properties: strain sweep of PIC hydrogels with different crosslinking approaches at 5 °C (A) and 37 °C (B). Conditions: For all samples: concentration of PIC = 2 mg mL$^{-1}$; the molar ratio of [linker]/[azide] is 0.4.

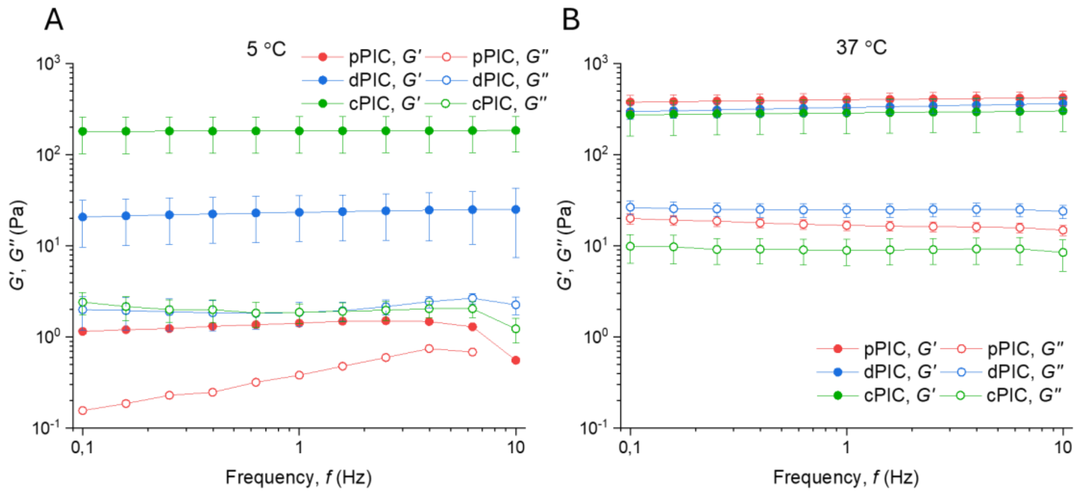


**Figure S5.** Viscoelastic properties: frequency sweep of PIC hydrogels with different crosslinking approaches at 5 °C (A) and 37 °C (B). Conditions: For all samples: concentration of PIC = 2 mg $mL^{-1}$; the molar ratio of [linker]/[azide] is 0.4.

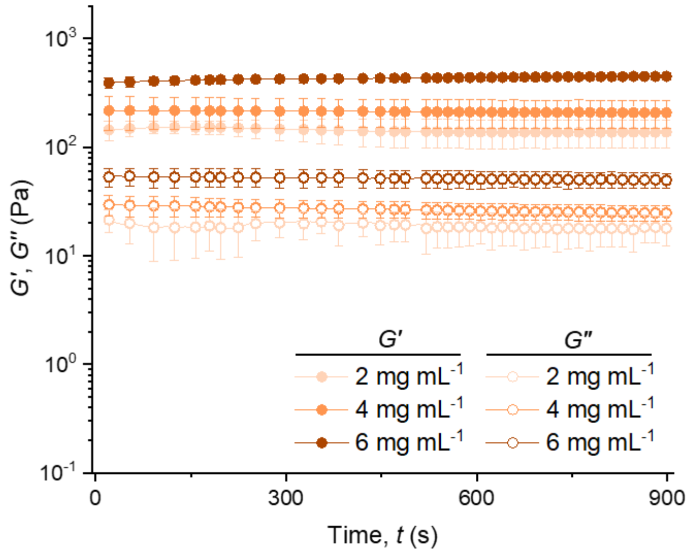


**Figure S6.** Time sweep of collagen hydrogels as a function of concentrations after gelation.

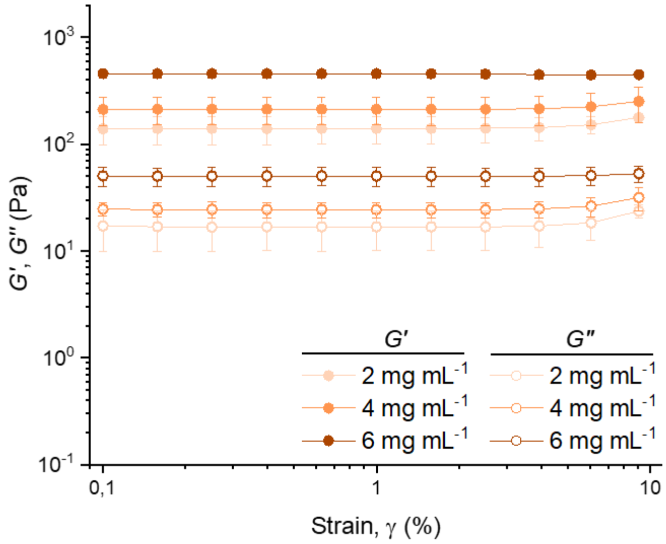


**Figure S7.** Viscoelastic properties: strain sweep of collagen gels as a function of polymer concentrations.

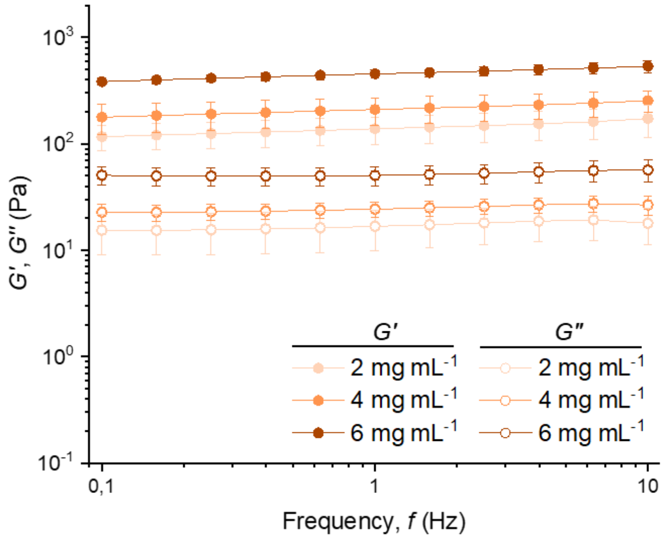


**Figure S8.** Viscoelastic properties: frequency sweep of collagen gels as a function of polymer concentrations.

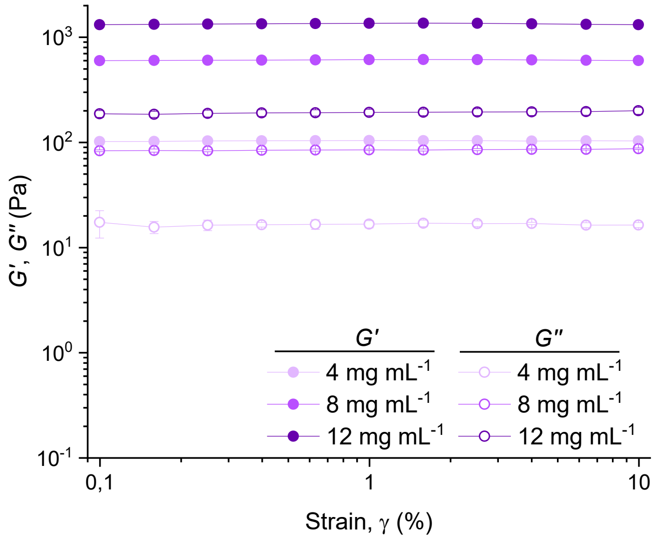


**Figure S9.** Viscoelastic properties: strain sweep of alginate hydrogels as a function of polymer concentrations.

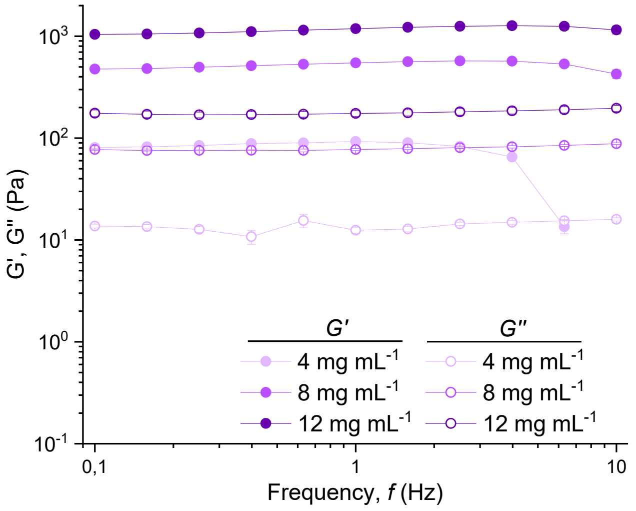


**Figure S10.** Viscoelastic properties: frequency sweep of alginate hydrogels as a function of polymer concentrations.

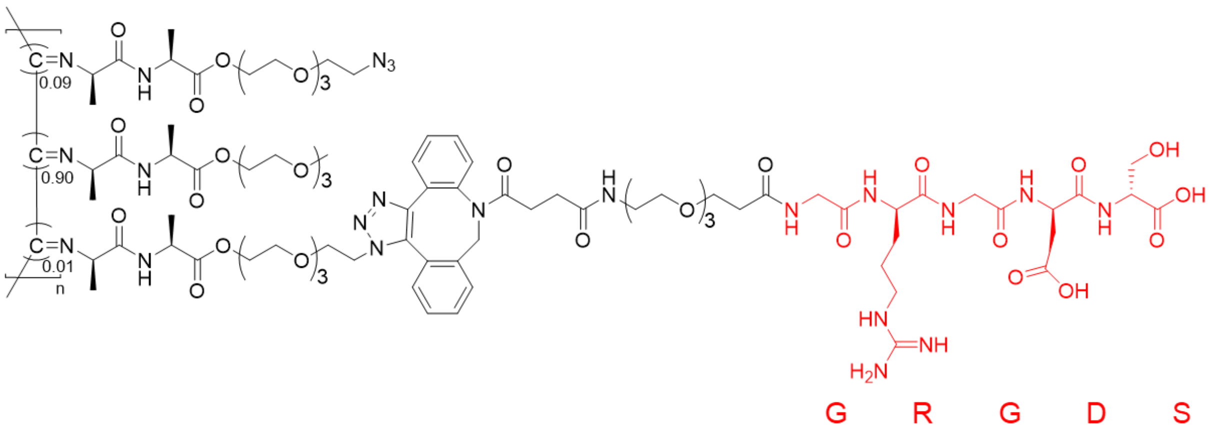


**Scheme S1.** Structure of RGD-functionalized polyisocyanides polymers.

**Table S1.** The crosslinking efficiency of different crosslinked PIC hydrogels.

| hydrogels | G′ at 5 °C (Pa) | G′ at 37 °C (Pa) | Crosslinking efficiency |
|---|---|---|---|
| pPIC | 1.4±0.2 | 370±60 | 0.004 |
| dPIC | 23±13 | 320±60 | 0.072 |
| cPIC | 179±75 | 290±120 | 0.617 |

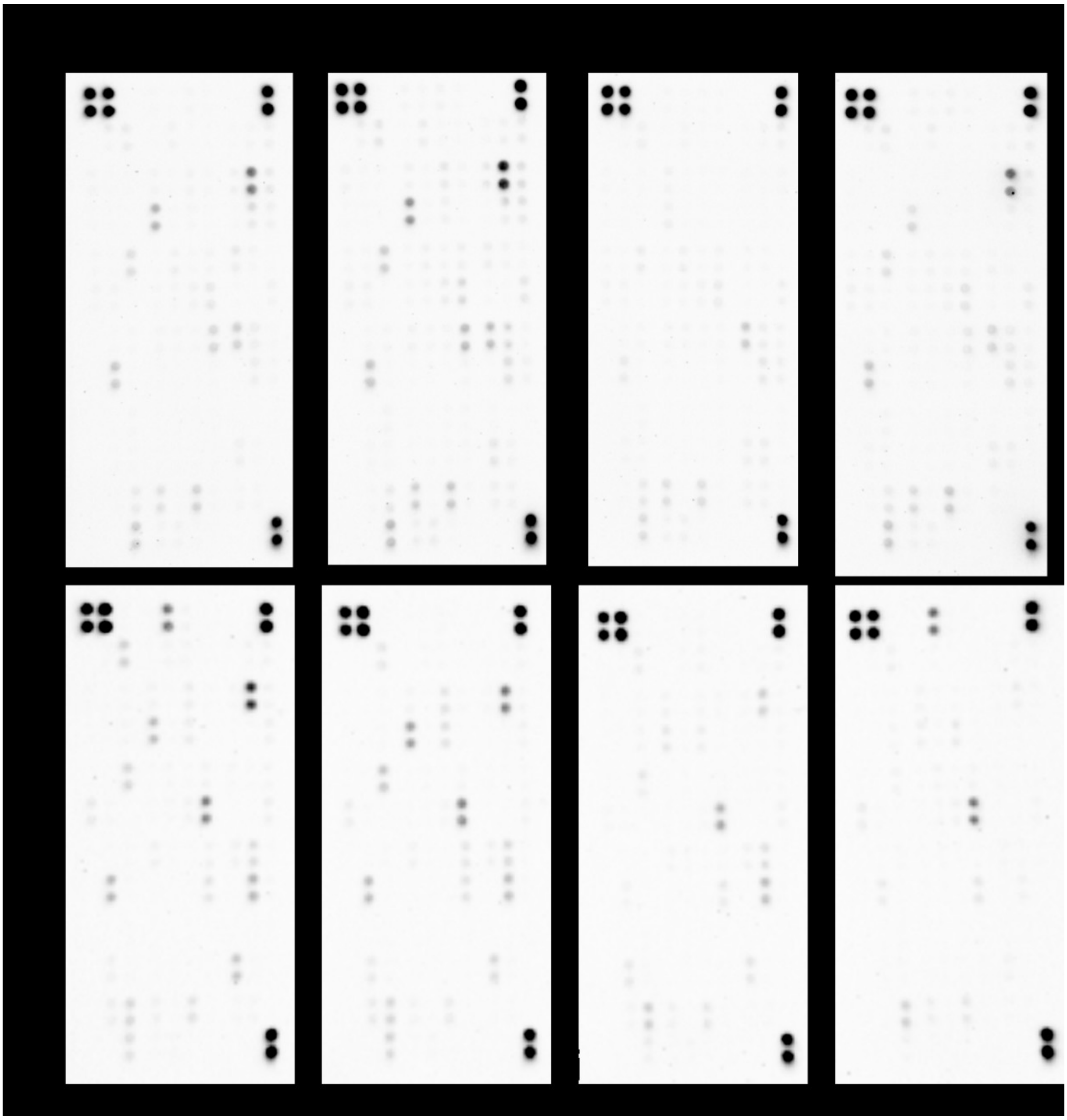

**Figure S11.** Chemiluminescence scanner images of cytokine secretion analysis from the initial experiment (A) and the corresponding replicate experiment (B). Conditioned media containing secreted cytokines were collected from hMSCs cultured in different hydrocups, including pPIC (i), dPIC (ii), collagen (iii), and alginate (iv), followed by incubation with antibody blot arrays targeting 105 different cytokines. Each antibody was represented by the pair of duplicate spots.

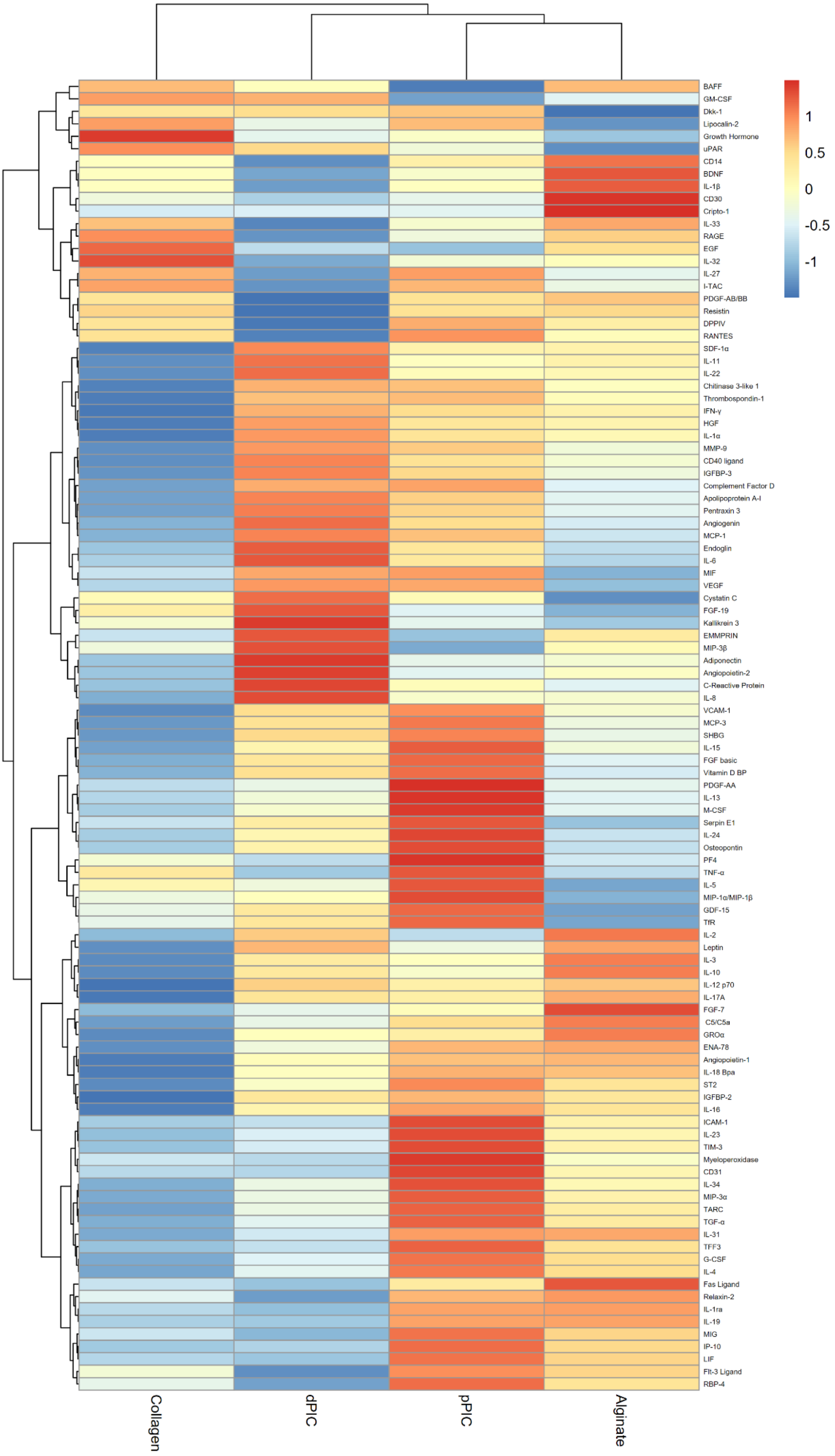

BAFF
GM-CSF
Dkk-1
Lipocalin-2
Growth Hormone
uPAR
CD14
BDNF
IL-1β
CD30
Cripto-1
IL-33
RAGE
EGF
IL-32
IL-27
I-TAC
PDGF-AB/BB
Resistin
DPPIV
RANTES
SDF-1α
IL-11
IL-22
Chitinase 3-like 1
Thrombospondin-1
IFN-γ
HGF
IL-1α
MMP-9
CD40 ligand
IGFBP-3
Complement Factor D
Apolipoprotein A-I
Pentraxin 3
Angiogenin
MCP-1
Endoglin
IL-6
MIF
VEGF
Cystatin C
FGF-19
Kallikrein 3
EMMPRIN
MIP-3β
Adiponectin
Angiopoietin-2
C-Reactive Protein
IL-8
VCAM-1
MCP-3
SHBG
IL-15
FGF basic
Vitamin D BP
PDGF-AA
IL-13
M-CSF
Serpin E1
IL-24
Osteopontin
PF4
TNF-α
IL-5
MIP-1α/MIP-1β
GDF-15
TfR
IL-2
Leptin
IL-3
IL-10
IL-12 p70
IL-17A
FGF-7
C5/C5a
GROα
ENA-78
Angiopoietin-1
IL-18 Bpa
ST2
IGFBP-2
IL-16
ICAM-1
IL-23
TIM-3
Myeloperoxidase
CD31
IL-34
MIP-3α
TARC
TGF-α
IL-31
TFF3
G-CSF
IL-4
Fas Ligand
Relaxin-2
IL-1ra
IL-19
MIG
IP-10
LIF
Flt-3 Ligand
RBP-4
1
0.5
0
-0.5
-1
Collagen
dPIC
pPIC
Alginate

**Figure S12.** Heatmap of cytokine profiles under different matrix conditions. Cytokine levels were row-scaled to visualize relative differences among collagen, dPIC, pPPIC, and alginate conditions, according to the chemiluminescence scanner images of cytokine secretion analysis shown in Figure S11. Red and blue colors indicate relatively higher and lower levels, respectively, for each cytokine. The displayed Z-score color scale is approximately −1.5 to +1.5. Four data sets were averaged.

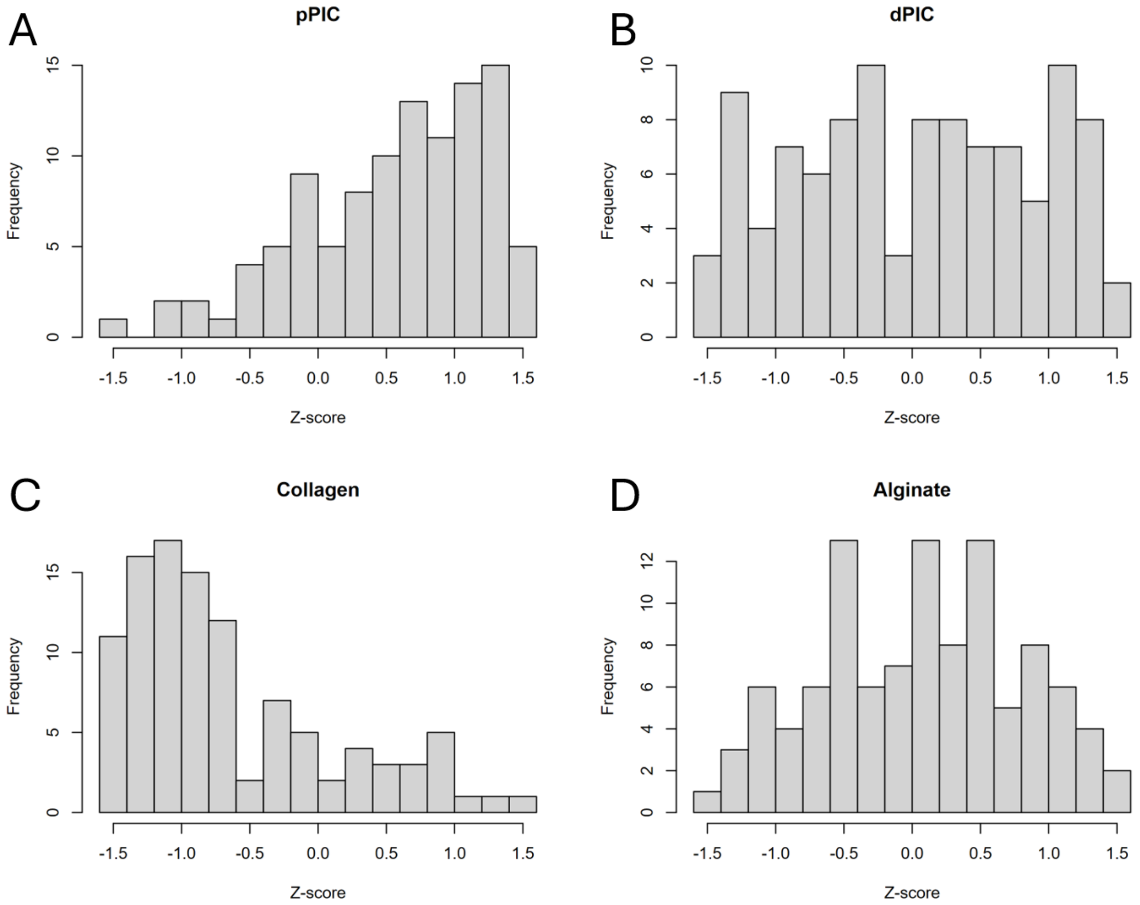


**Figure S13.** Histogram analysis of cytokine intensity values and relative frequencies for each of the hydrogel conditions included in the heatmap cluster analysis in Figure S12.

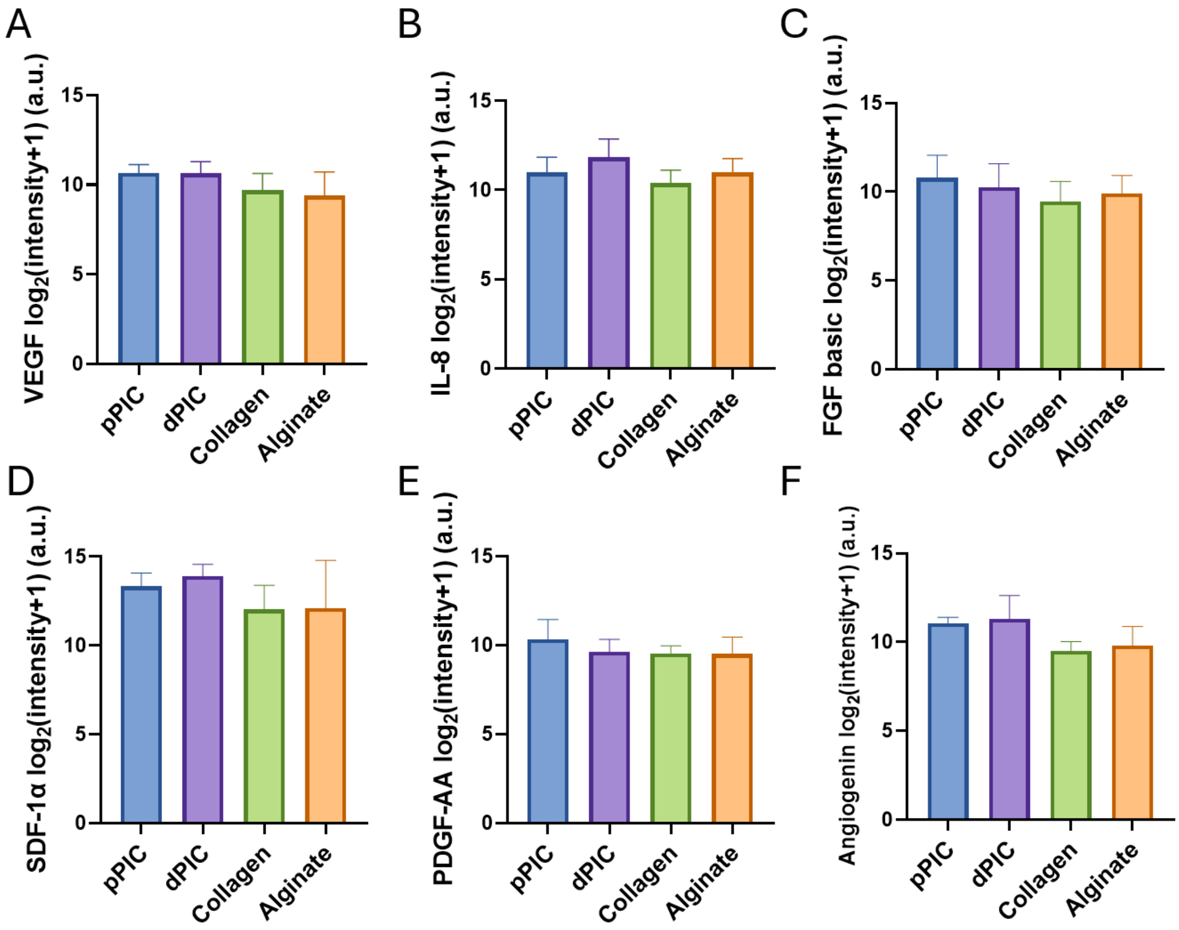


**Figure S14.** Intensity of the expressed angiogenesis-related cytokines, including VEGF (A), IL-8 (B), FGF basic (C), SDF-1α (D), PDGF-AA (E), and angiogenin (F). Data were tested through a one-way ANOVA, followed by the Tukey's multiple comparison tests. Probability values $p > 0.05$ were considered not significant. There was no significant difference.

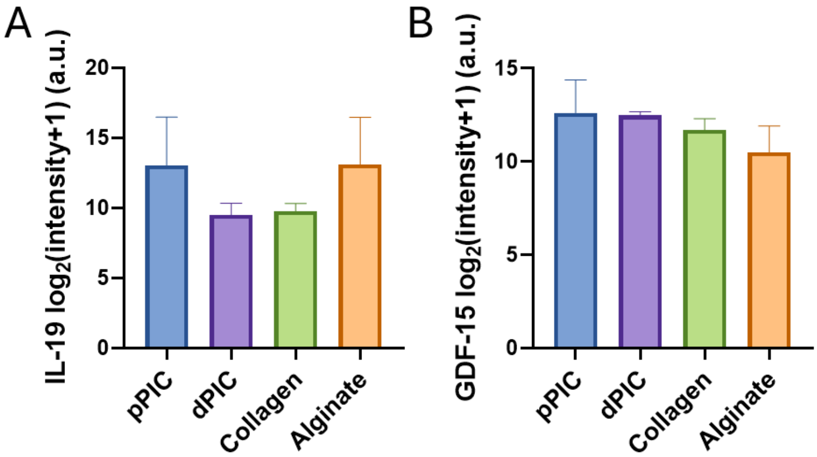


**Figure S15.** Intensity of the expressed immunomodulatory-related cytokines, including IL-19 (A), and GDF-15 (B). Data were tested through a one-way ANOVA, followed by the Tukey's multiple comparison tests. Probability values $p > 0.05$ were considered not significant. There was no significant difference.

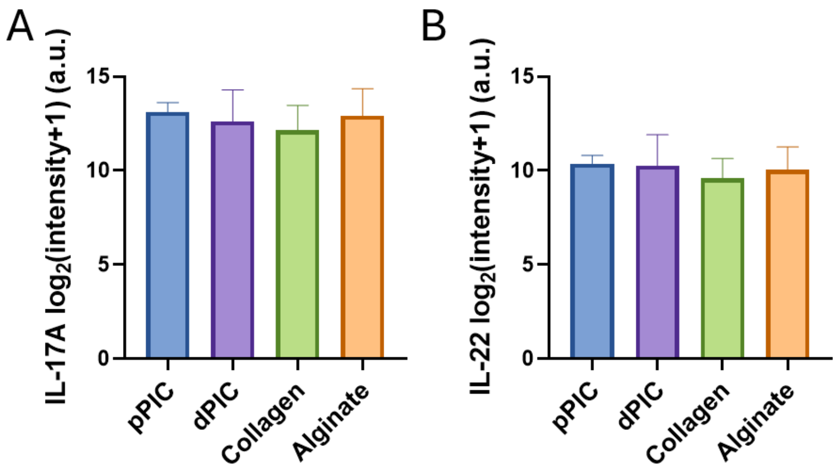


**Figure S16.** Intensity of the expressed migratory-related cytokines, including IL-17A (A) and IL-22 (B). Data were tested through a one-way ANOVA, followed by the Tukey's multiple comparison tests. Probability values $p > 0.05$ were considered not significant. There was no significant difference.

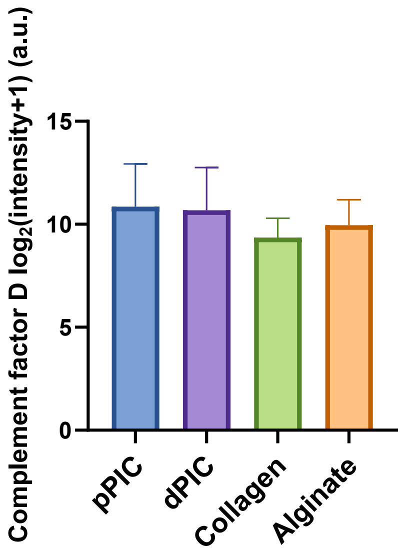


**Figure S17.** Intensity of the expressed complement factor D cytokines. Data were tested through a one-way ANOVA, followed by the Tukey's multiple comparison tests. Probability values $p > 0.05$ were considered not significant. There was no significant difference.
